# Ultradeep optical coherence tomography

**DOI:** 10.64898/2026.10.28.762671

**Authors:** Santosh Balakrishnan, Gauri Arora, Steven G. Adie

## Abstract

Deep optical imaging of intact brain tissue is limited by scattering and absorption, restricting our ability to study subcortical structures. Here, we introduce ultradeep optical coherence tomography (OCT), which combines hardware and computational approaches to enhance the signal from deep tissue regions. Ultradeep OCT utilizes a longer illumination center wavelength of 1575 nm, spectral-domain balanced detection and imaging in the conjugate configuration with the zero optical path difference positioned deep in the tissue. Autocorrelation artifacts are measured and subtracted, and multiple frames are coherently averaged to improve the signal-to-noise ratio further. These techniques, along with computational refocusing and bandwidth control, allowed us to extend structural imaging depth in biological tissues. Ultradeep OCT demonstrated physical imaging depths of 2.3 to 2.8 mm (mean 2.5 mm) in the *ex vivo* mouse brain and 2.5 to 3.0 mm (mean 2.8 mm) in chicken tissue. The resulting volumes in the *ex vivo* mouse brain samples depict the dorsal hippocampus and its two major subregions with sufficient contrast to allow segmentation and three-dimensional reconstructions. Features in the thalamus could also be discerned. These findings demonstrate the potential of ultradeep OCT as a label-free approach for visualizing deep-brain structures at a spatial scale complementary to magnetic resonance imaging and microscopy. This approach could enable optical imaging studies of hippocampal structural changes in mouse models of neurodegenerative disease.

## Introduction

Structural changes associated with tissue development, remodeling, and pathology can extend beyond the superficial depths that are typically accessible to optical imaging. Visualizing such changes in intact biological specimens, particularly longitudinally, therefore requires methods capable of imaging at depth while preserving tissue integrity. Label-free optical imaging can provide depth-resolved views of tissue morphology, but scattering reduces visibility of tissue structures as depth increases [Dunsby, 2003; Yoon, 2020]. In the brain, extending imaging depth could provide access to deep cortical and subcortical structures that are crucial for cognition and are affected by neurodegenerative disease [de Flores, 2015]. At these depths, scattering and absorption significantly attenuate backscattered light, reducing the signal-to-noise ratio (SNR) [Dunsby, 2003; Yoon, 2020]. Optical coherence tomography (OCT) is a promising approach for label-free imaging in scattering tissue because the combination of confocal and coherence gating allows selective detection of backscattered light from deeper tissue regions [Dunsby, 2003; Bouma, 2022]. Nevertheless, with increasing propagation distance, attenuation reduces the ballistic and quasi-ballistic light available for image formation, while the contribution from multiple scattering also increases [Yoon, 2020]. These phenomena impose a tissue-dependent limit on achievable OCT imaging depth.

Optical coherence microscopy (OCM) has enabled high-resolution *ex vivo* [Bolmont, 2012; Assayag, 2013; Min, 2015; Yamanaka, 2019] imaging of human or rodent brains and *in vivo* [Bolmont, 2012; Srinivasan, 2012; Leahy, 2013; Zhu, 2021] imaging of rodent brains. Despite this progress, label-free visualization of neurites, myelin fibers or markers of neurodegenerative diseases, including amyloid-β plaques, neurofibrillary tangles, or other microstructural changes, remains challenging at depth [Magnain, 2019; Gesperger, 2019; Zhu, 2021]. Structural OCT addresses a complementary imaging regime by prioritizing penetration depth and tissue-scale architecture at modest spatial resolution, typically 10 to 20 µm [Roper, 1998; Choi, 2015; Chong, 2015]. Imaging performance for structural imaging is strongly influenced by the system dynamic range (DR) and the SNR at depth. Deep structural OCT of the brain has mostly been used to demonstrate penetration depth or to support angiography [Choi, 2015; Chong, 2015; Chen, 2016; Park, 2018], rather than to analyze tissue architecture; neurodegeneration-related changes have instead been studied mainly with high-resolution OCM [Bolmont, 2012; Zhu, 2021].

Here, we introduce ultradeep OCT, a combination of hardware, acquisition, and computational techniques designed to extend structural OCT imaging depth. Ultradeep OCT combines a longer center wavelength of 1575 nm, balanced detection spectral-domain OCT (SD-OCT) [Bradu, 2012], conjugate imaging with the zero optical path difference (OPD) positioned deep in the tissue [Spaide, 2008], measurement and subtraction of autocorrelation (AC) artifacts [Ai, 2006], repeated acquisitions with coherent averaging [Tomlins, 2007], and computational refocusing [Adie, 2012]. This combination of techniques extends the depth range that is supported by the DR of the imaging system via a cumulative improvement of SNR. The impact of multiple scattering is also reduced by using the longer center wavelength.

Using ultradeep OCT, we demonstrate a mean physical structural imaging depth of 2.5 mm (range 2.34 to 2.79 mm, five measurements) in the *ex vivo* mouse brain and 2.77 mm (range 2.49 to 3.0 mm, three measurements) in chicken tissue, with physical depth calculated using a refractive index of 1.35. Increased imaging depth in the *ex vivo* mouse brain with the 1575 nm system compared with 1300 nm is also demonstrated. To compare structural imaging depth quantitatively across techniques, a signal-to-background ratio (SBR) metric as a function of depth was calculated. The maximum structural imaging depth was defined as the depth at which the SBR fell below a user-specified threshold. This approach reduces some reliance on mean intensity profiles or subjective assessments of feature visibility used in previous studies.

Previous *in vivo* studies have reported structural imaging depths of up to 2.1 mm at 1700 nm through a reinforced thinned-skull window [Chong, 2015], 1.9 mm at 1700 nm through a glass cranial window [Park, 2018], and 2.3 mm at 1300 nm through a glass cranial window [Choi, 2015]. High-resolution imaging to approximately 1.6 mm has also been demonstrated at 1700 nm through a reinforced thinned-skull window [Zhu, 2021]. In the *ex vivo* mouse brain, the structural imaging depth achieved here exceeds previously reported values at any wavelength for fresh, uncleared tissue [Bolmont, 2012; Leahy, 2013; Yamanaka, 2016; Yamanaka, 2019], to the best of our knowledge. Imaging at 1575 nm also permits the use of standard InGaAs detectors, which can provide higher quantum efficiency than the extended InGaAs detectors required for operation in the 1700 nm spectral band [Chong, 2015; Zhu, 2021].

Notably, ultradeep OCT enabled visualization of the dorsal hippocampus and its major subregions, the cornu ammonis (CA) and dentate gyrus (DG), within the intact *ex vivo* mouse brain, with sufficient contrast and spatial resolution to enable segmentation and three-dimensional volumetric reconstructions. Hippocampal subfield atrophy, assessed using magnetic resonance imaging (MRI), has been shown to be a sensitive biomarker of neurodegenerative disease in humans [de Flores, 2015; Bouwman, 2025]. Although evidence for hippocampal volume loss in mouse models of Alzheimer’s disease is mixed, MRI studies have reported disease-associated reductions in regions of the hippocampal formation in some models [Jullienne, 2022]. Ultradeep OCT could bridge the gap between MRI and high-resolution optical microscopy, providing label-free visualization of hippocampal anatomy that may support quantitative studies of deep-brain structural changes in mouse models of neurodegeneration.

## Results

### Complementary hardware and computational advances enable ultradeep OCT

Ultradeep OCT is a combination of hardware, acquisition, and computational strategies that improve signal detection and suppress noise to cumulatively improve imaging depth (Fig. 1(a)). Conventional OCT, defined here as imaging at 1300 nm in the standard configuration with a single spectrometer, provided a maximum structural imaging depth of 1.40 mm in the representative *ex vivo* mouse brain result shown in Fig. 1(b), 1(c). Maximum imaging depth was defined as the depth at which the SBR fell below 7.04 dB for this dataset.

**Figure 1.**
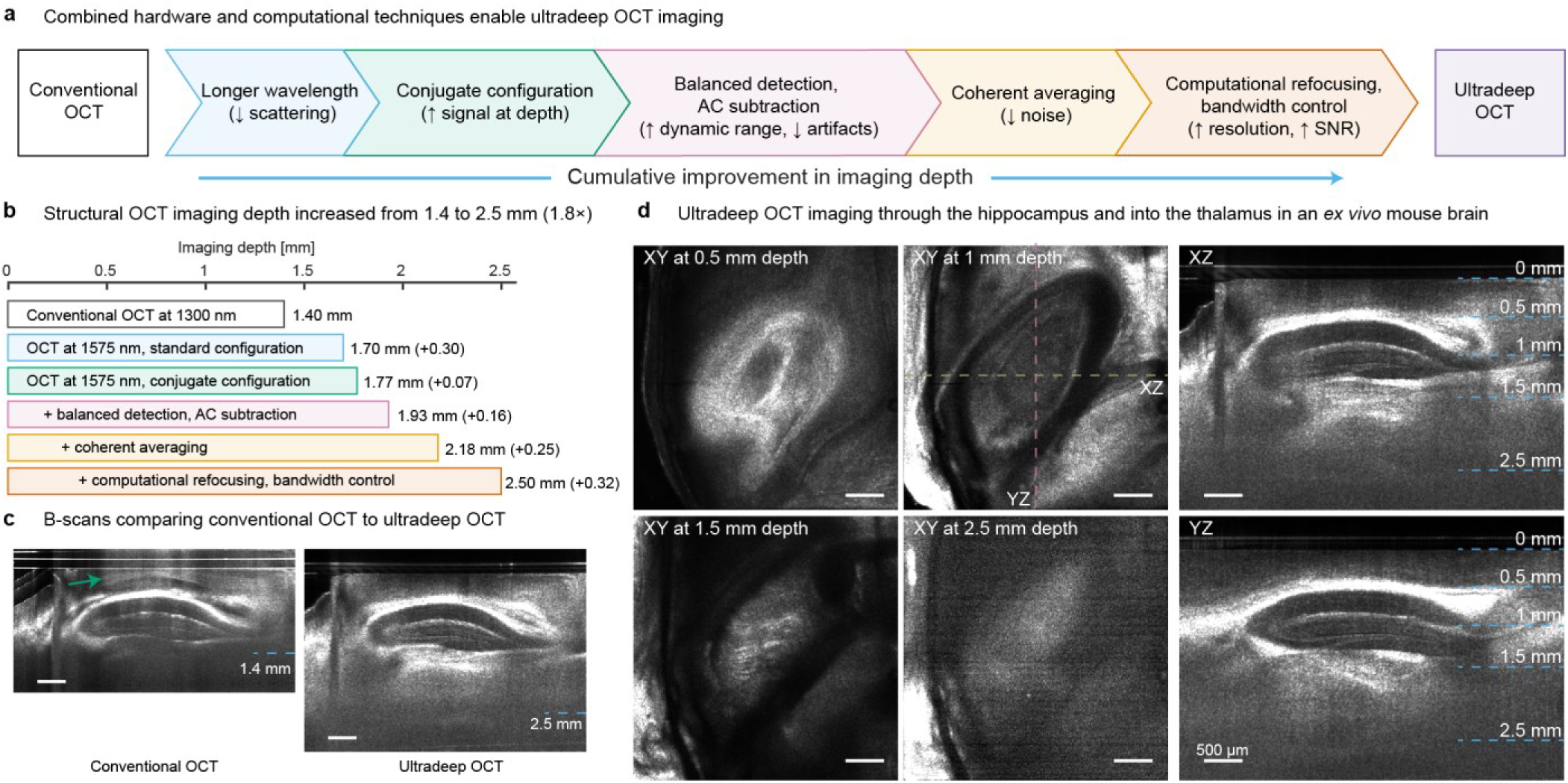
Combination of hardware and computational techniques enables ultradeep OCT. **a,** Overview of the techniques in ultradeep OCT. AC, autocorrelation; SNR, signal-to-noise ratio. **b,** Structural imaging depth in one representative *ex vivo* mouse brain dataset after implementation of each technique. Values next to the bars indicate maximum imaging depth; values in parentheses indicate the observed change associated with each successive technique relative to the preceding system configuration. Maximum depth was defined by the signal-to-background ratio threshold method described in Methods. **c**, Representative XZ B-scans of the same *ex vivo* mouse brain acquired using conventional OCT and ultradeep OCT. Dashed blue lines indicate maximum imaging depths. The arrow on conventional OCT indicates AC artifacts. **d**, XY, XZ, and YZ views from the ultradeep OCT volume. XY images are shown at the indicated physical depths. Dashed lines in the 1 mm XY image identify the locations of the XZ and YZ views. Cross-sectional images are shown after intensity normalization in depth. Contrast was set independently for each image. Scale bars, 500 µm.

Imaging with the 1575 nm system in the standard configuration extended the imaging depth to 1.70 mm (Fig. 1(b)), while imaging at 1575 nm in the conjugate configuration, which improves signal collection from deep tissue regions, further increased the imaging depth to 1.77 mm. Balanced detection combined with autocorrelation (AC) subtraction increased the dynamic range, suppressed noise and removed autocorrelation artifacts, extending the imaging depth to 1.93 mm. Coherent averaging subsequently increased the imaging depth to 2.18 mm by reducing the noise floor at depth. Finally, computational refocusing with bandwidth control improved resolution away from the focal plane, while also improving the SNR at depth, resulting in a maximum structural imaging depth of 2.50 mm in this *ex vivo* mouse brain sample. These advances increased the imaging depth by 1.10 mm, corresponding to a 1.8-fold improvement as compared to conventional OCT (Fig. 1(b)).

Volumetric ultradeep OCT imaging allowed visualization through the dorsal hippocampus and into the thalamus of the *ex vivo* mouse brain (Fig. 1(c), 1(d)). Representative cross-sectional images, shown in Fig. 1(c), illustrate deeper imaging with ultradeep OCT as compared with conventional OCT. The bright horizontal lines above the tissue surface are from a coverslip that was placed on the sample surface during imaging. In addition to extending the detectable depth, the improved visualization of the dorsal hippocampal features was evident in the ultradeep results. Deeper regions in the thalamus that were not discernible using conventional OCT could be visualized with ultradeep OCT.

*En face* images acquired at progressively greater depths (Fig. 1(d)) revealed distinct structural features throughout the volume. At 0.5 mm beneath the brain surface, the images showed features consistent with the corpus callosum (CC). At 1 mm, laminar structures similar to the stratum lacunosum-moleculare (SLM) of the CA were visible. Structures identified as probable myelinated tracts in the thalamus were detected on the XY image at 1.5 mm depth. Even at a depth of 2.5 mm, which represents regions in the deep thalamus, while specific features could not be identified, some structural contrast remained detectable. The corresponding orthogonal XZ and YZ views depicted the dorsal hippocampus across its full depth and demonstrated continuous structural imaging from the tissue surface to regions of the thalamus. Across available measurements in *ex vivo* brains, the structural imaging depth increased from a mean value of 1.53 mm (n = 2) for conventional OCT to 2.53 ± 0.18 mm (mean ± s.d., n = 5) for ultradeep OCT. See S4 for additional details.

### Improved structural visibility at depth with the 1575 nm system

To compare the system-level penetration depth in the two wavelength bands, OCT volumes acquired from the same *ex vivo* mouse brain at 1300 and 1575 nm were compared (Fig. 2). Both datasets were acquired in the standard imaging configuration with balanced detection, over similar fields of view (FOVs), and with identical camera line rates and exposure times, but with system-specific optical powers, imaging ranges and sensitivities.

**Figure 2.**
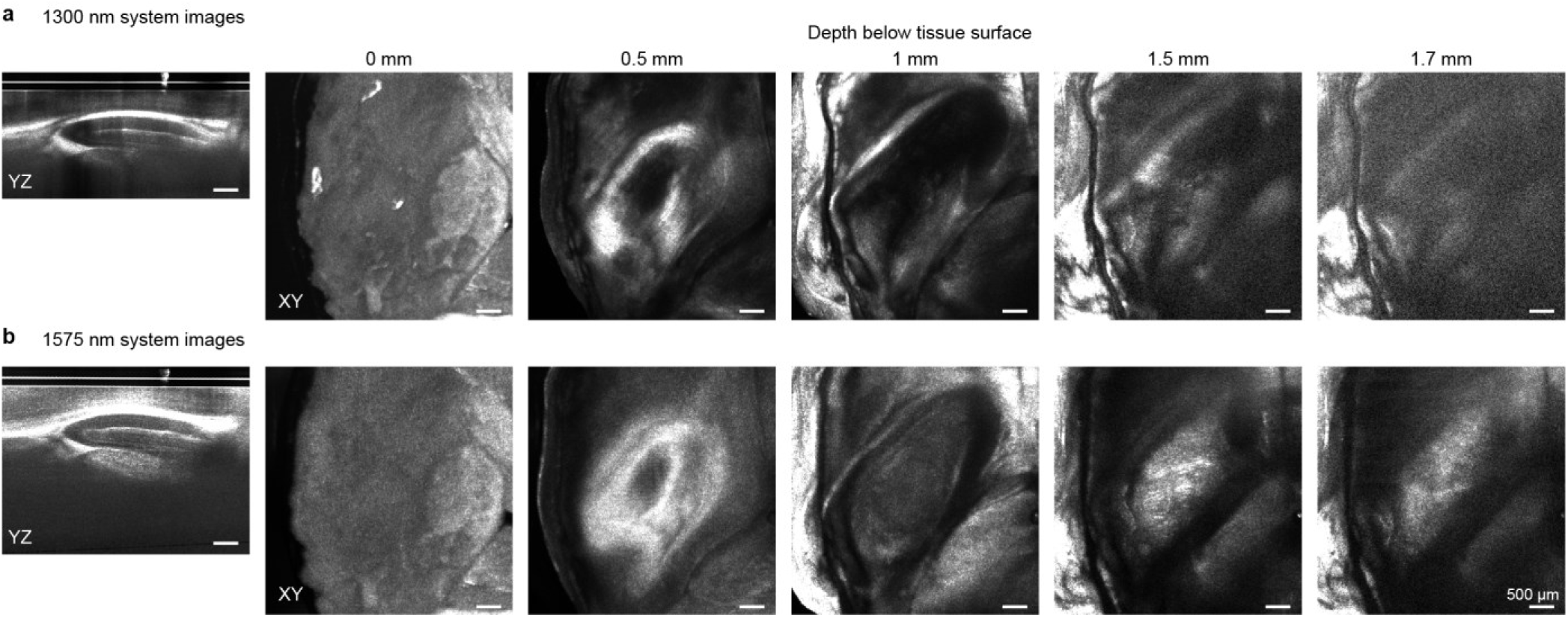
Enhanced structural visibility at depth in the *ex vivo* mouse brain for the 1575 nm system as compared to the 1300 nm system. **a,** YZ cross-sectional image, and XY *en face* images at depths of 0, 0.5, 1, 1.5 and 1.7 mm below the tissue surface, acquired at 1300 nm in the standard imaging configuration with balanced detection. **b,** Corresponding images of the same sample, acquired over a similar FOV at 1575 nm, standard configuration with balanced detection. The contrast of each image was adjusted independently. Scale bars, 500 µm.

At shallow depths, both wavelengths resolved comparable features within the tissue (Fig. 2(a), 2(b)). The *en face* images acquired at the tissue surface and 0.5 mm below the surface revealed similar gross tissue organization, although the relative intensities of individual structures differed between wavelengths. The benefit of imaging with the 1575 nm system becomes increasingly apparent with depth. At 1 mm below the tissue surface, layered anatomical features remained visible at both wavelengths, but the 1575 nm image showed better structural contrast, enabling identification of hippocampal layers. At 1.5 mm, structural contrast was substantially reduced in the 1300 nm dataset, whereas the 1575 nm result showed bright features within the thalamus. At 1.7 mm, the 1300 nm image was dominated by background with limited discernible tissue structure. In contrast, the corresponding 1575 nm image retained localized structural contrast and resolved features that were not readily visible at 1300 nm (Fig. 2(a), 2(b)).

The corresponding cross-sectional YZ images further demonstrated the wavelength-dependent difference in penetration. Although both datasets visualized superficial and intermediate-depth structures, signal attenuation with depth was more pronounced at 1300 nm. Imaging at 1575 nm yielded better structural contrast over a greater depth, providing improved visualization of the deeper brain regions. See S1 and S5 for additional details.

### Conjugate configuration extends imaging depth

We next investigated whether the depth-dependent sensitivity roll-off of the OCT system, which is typically viewed as a detriment for deeper imaging, could advantageously be used to improve signal detection from deeper tissue. In the standard imaging configuration, signals from increasing depths are subject to both the tissue-induced attenuation and the spectrometer roll-off, as shown in the simulation in Fig. 3(a). The combined effects of these two mechanisms cause the detected signal to decrease more rapidly with depth than would be expected from tissue attenuation alone. The maximum imaging depth, *d_max_*_,*std*_, is reached when the attenuated signal falls below the minimum detectable signal determined by the system DR.

**Figure 3.**
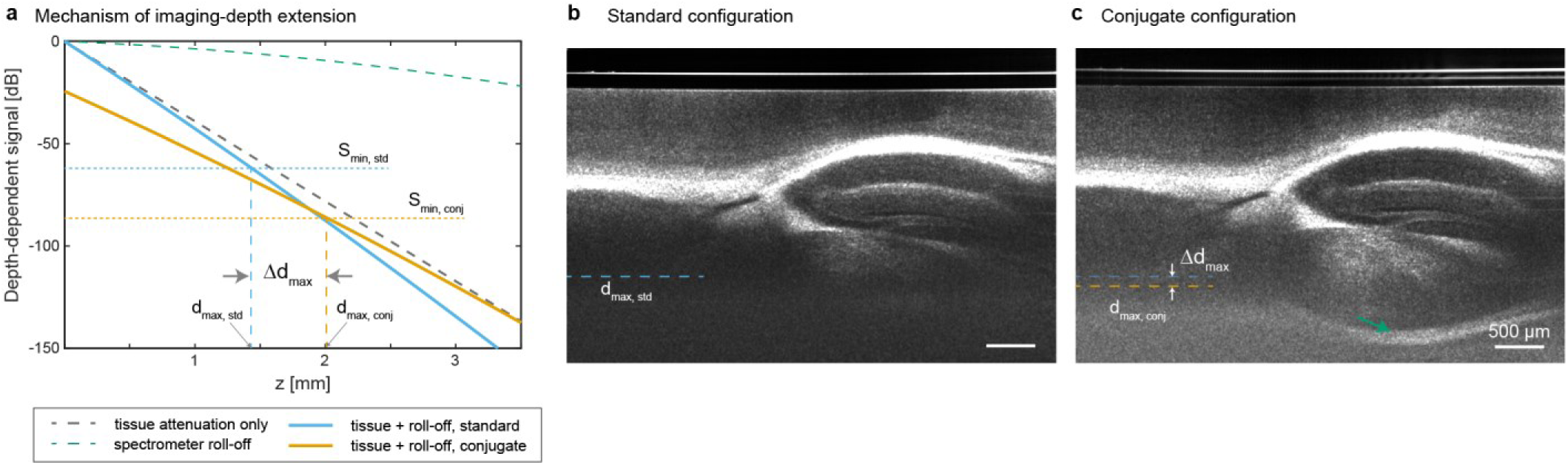
Extension of imaging depth in the conjugate configuration. **a,** Simulated attenuation curves showing the combined effects of tissue attenuation and spectrometer roll-off. In the standard configuration, roll-off increases the attenuation of signals originating from deeper tissue, limiting the detectable depth to *d*max,std. In the conjugate configuration, the roll-off contribution is reversed with respect to the imaging depth, extending the detectable depth to *d*max,conj. The resulting depth gain is denoted by Δ*d*max. The horizontal dashed lines represent the minimum detectable signal level given by *Smin* = *S*(0) − *DR*, where *DR* is the dynamic range. **b,c,** Representative B-scans of *ex vivo* mouse brain acquired at 1575 nm with balanced detection using the standard (**b**) and conjugate (**c**) configurations. B-scans are shown without intensity normalization, and image contrast was set independently. Dashed lines mark the maximum imaging depths and the gain in imaging depth is denoted by Δ*dmax*. The arrow in (c) indicates AC artifacts. Scale bars, 500 µm.

In the conjugate imaging configuration, the zero OPD position is placed deep in the sample. Consequently, deeper tissue signals occur closer to the region with lower spectrometer roll-off, reversing the contribution of roll-off with respect to tissue depth (Fig. 3(a)). This arrangement reduces sensitivity for shallower regions in the sample as these now experience more spectrometer roll-off attenuation, but the sensitivity at greater depths is improved in exchange. Therefore, for a given system DR, a greater imaging depth, *d_max_*_,*conj*_, is obtained, corresponding to an increase of Δ*d_max_* compared to the standard configuration (Fig. 3(a)). The simulation assumes homogeneous brain tissue with an attenuation coefficient of 4.5 mm^−1^, system dynamic range of 62 dB and 6 dB roll-off distance of 1.46 mm in tissue. The zero OPD for the conjugate configuration was placed at *z* = 3.5 mm. In Fig. 3(a), *S_min_*_,*std*_ and *S_min_*_,*conj*_ are the minimum detectable signal levels in the standard and conjugate configurations, respectively. Note that the system noise floor does not change between the standard and conjugate configurations. The minimum detectable signal levels, *S_min_*_,*std*_ and *S_min_*_,*conj*_, are shown offset from the signal level at zero depth, *S*(0), by an amount equal to the system DR to allow comparison between the two configurations. For a given system DR, this means that when the upper limit of the DR is set closely above the maximum backscattered signal level from the sample, the conjugate acquisition approach allows the minimum detectable lower limit to correspond to a greater imaging depth. These results, which are meant to be illustrative, depict an approximate increase in imaging depth of 0.55 mm and a 2 mm maximum imaging depth with the conjugate configuration.

The benefit of imaging in the conjugate configuration is illustrated in Fig. 3(b) and 3(c) by imaging the same region of an *ex vivo* mouse brain using balanced detection at 1575 nm with the standard and conjugate configurations. Both datasets were acquired with identical settings, except for reference power adjustment when switching from the standard to the conjugate imaging configuration. In the standard configuration, tissue structure was visible through the cortex and portions of the dorsal hippocampus, but the signal decreased past the hippocampus and became indistinguishable from the background (Fig. 3(b)). In the conjugate configuration, the ventral portions of the hippocampus were better resolved and signal from the thalamus was also evident (Fig. 3(c)). The conjugate imaging B-scan also exhibits AC artifacts (arrow in Fig. 3(c)) that are present even with balanced detection.

### Balanced detection with AC subtraction reveals deep tissue structures

Conjugate imaging improved sensitivity to signals from deeper tissue but also caused AC artifacts, generated by strongly scattering interfaces, to overlap with the deep tissue signal. In the *ex vivo* mouse brain, strong reflections from the coverslip and highly scattering white-matter structures, including the CC, produced prominent AC artifacts at larger depths (Fig. 4(a), magenta arrows). These artifacts limited the visibility of deep structures despite the improved depth-dependent sensitivity provided by the conjugate configuration.

**Figure 4.**
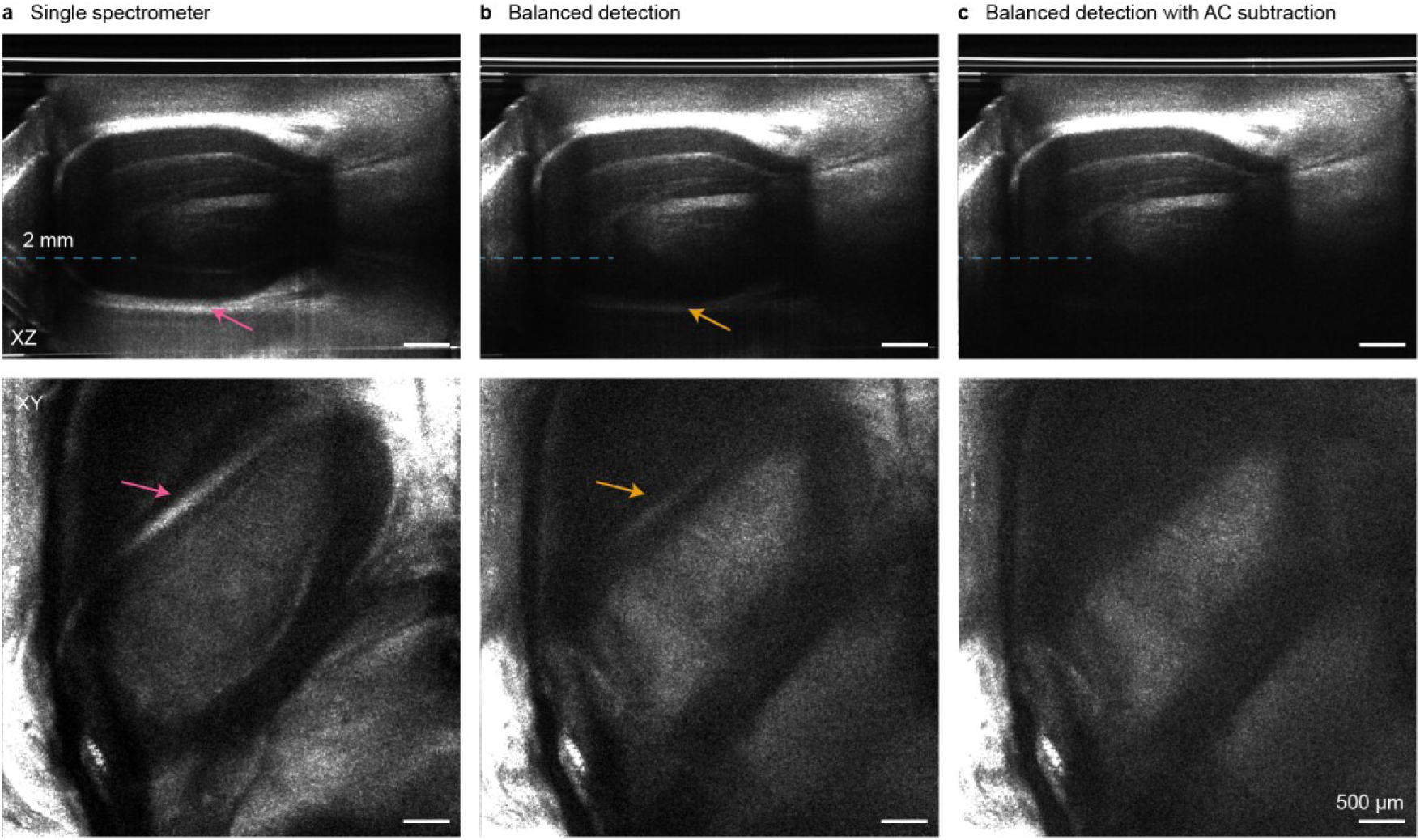
Balanced detection with AC subtraction suppresses artifacts that obscure deep brain features. Representative XZ and XY images in an *ex vivo* mouse brain acquired at 1575 nm using the conjugate configuration. **a,** Images for single-spectrometer detection. **b,** Balanced detection. **c,** Balanced detection with AC subtraction. Dashed blue lines on the XZ images indicate the depth of the corresponding *en face* XY images. Magenta arrows identify AC artifacts on the single-spectrometer B-scans; orange arrows mark the residual artifacts after balanced detection alone. Images in each row are shown with the same display settings. XZ images shown without depth-dependent intensity normalization. Scale bars, 500 µm.

Balanced detection, which makes use of the detected signals from two spectrometers, was implemented as it not only suppresses the common-mode AC artifacts and source relative intensity noise (RIN) but also doubles the useful interferometric signal [Balakrishnan, 2026]. The results shown in Fig. 4(b) indicate that balanced detection markedly reduced AC artifacts allowing additional deep tissue features to become visible. However, residual AC artifacts (Fig. 4(b), orange arrows) that resembled superficial tissue structures, remained due to the amount of common-mode rejection achievable [Balakrishnan, 2026]. Owing to the lower SNR for the tissue features at larger depths, even weak residual AC artifacts after balanced detection obscured underlying structural information.

An additional step of AC subtraction was introduced to further reduce AC artifact visibility. The AC artifacts were first measured by blocking the reference arm and then subtracted from the spectrometer signal before spectrometer matching and balanced signal calculation. The reduction in AC artifacts is apparent in both the cross-sectional and *en face* images after balanced detection with AC subtraction shown in Fig. 4(c). The *en face* image also illustrates that the combination of improved SNR from balanced detection along with the additional AC subtraction step improves visualization of tissue features at 2 mm depth. See S3 for additional information.

### Coherent averaging lowers the noise floor but is limited by signal decorrelation

Coherent averaging was used as a means of reducing the noise floor and extending the detectable imaging depth. Because coherent averaging preserves systematic signals while suppressing uncorrelated noise, theoretically the SNR is expected to increase in proportion to √*n*, where *n* is the number of frames averaged [Tomlins, 2007; Pfeiffer, 2020]. The phase of a coverslip, placed on top of the sample, was registered across frames before coherent averaging to correct for any system instabilities [Liu, 2018; Leartprapun, 2021]. However, the improvement in SNR requires that the tissue signal remains correlated throughout the acquisition.

In chicken tissue, coherent averaging of 16 consecutively acquired frames reduced the background and improved the visibility of weak signals at depth (Fig. 5(a)). Compared with the single B-scan, the coherently averaged cross-sectional image exhibited a lower noise floor, which improved the contrast for features deeper in the tissue. Mean A-line profiles obtained from the B-scans are shown in Fig. 5(b). The plots demonstrate that the coherent averaging lowered the noise floor at depth by approximately 4 dB. This reduction in the noise floor enabled the visualization of the feature at approximately 3.25 mm below the tissue surface, as indicated by the green arrow in Fig. 5(b).

**Figure 5.**
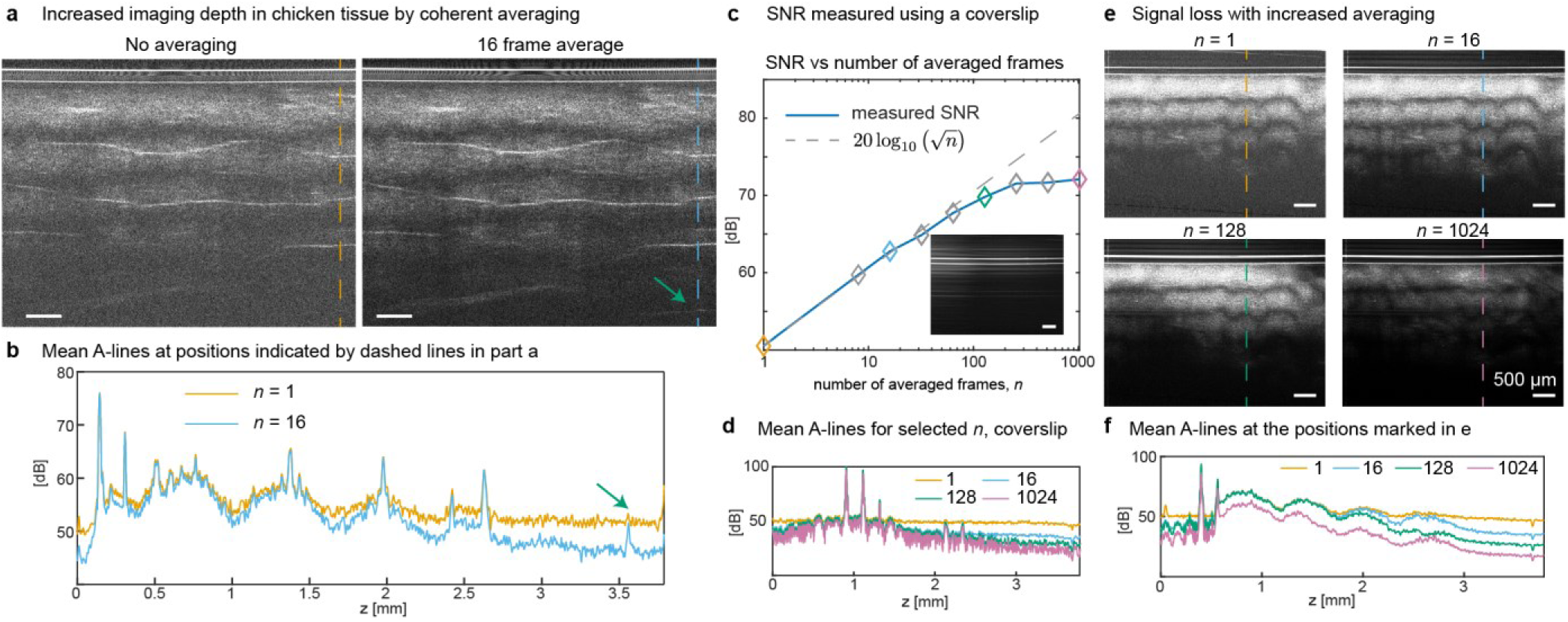
Coherent averaging lowers the noise floor and increases the imaging depth until signal decorrelation limits performance. **a,b,** Increased imaging depth in chicken tissue with 16-frame coherent averaging. **a,** Representative B-scans without averaging and with 16-frame coherent averaging. **b,** Mean of 64 A-lines centered at the positions marked by the dashed lines in **a**. The green arrow marks a weak tissue feature at 3.25 mm from the top of the tissue that becomes visible after coherent averaging. **c,** Coverslip SNR as a function of the number of coherently averaged frames, *n*, compared with the theoretical √*n* scaling. The inset shows a representative B-scan of the coverslip. **d,** Mean A-lines for selected values of *n* in **c**. **e,f,** Signal loss with prolonged averaging caused by decorrelation. **e,** Representative B-scans of chicken tissue with 1, 16, 128, or 1024 frame coherent averaging. **f,** Mean of 64 A-lines at the positions marked by the dashed lines in **e**. B-scans within each comparison are shown with identical contrast settings and without depth-dependent intensity normalization. Scale bars, 500 µm.

To characterize coherent averaging performance under stationary conditions, SNR as a function of *n* was measured using a coverslip (Fig. 5(c)). SNR was calculated as the ratio of the coverslip peak amplitude to the mean noise amplitude for z from 3 to 3.5 mm, *SNR* [*dB*] = 20 log_10_(*s_peak_*/*μ_noise_*). The SNR initially followed the theoretical 20 log_10_ √*n* dependence, increasing from approximately 50 dB without averaging to approximately 65 dB at intermediate averaging levels. At larger values of *n*, however, the measured SNR deviated from ideal scaling and approached a plateau of approximately 72 dB, with limited additional improvement beyond approximately 128 frames. The corresponding mean A-line profiles showed a progressive reduction in the noise floor while the coverslip signal remained approximately constant (Fig. 5(d)). These measurements indicated that coherent averaging produced the expected SNR benefit over a range of averaged frames, after which residual correlated noise limited further improvement.

The effect of coherently averaging an increasing number of frames was also investigated with chicken tissue (Fig. 5(e), 5(f)). Increasing the number of coherently averaged frames from 1 to 16 lowered the background while largely preserving tissue contrast. However, with prolonged averaging, tissue features were lost owing to signal decorrelation, most likely caused by sample motion, as seen on the 128- and 1024-frame averaged results. The signal loss was substantially greater at 1024 frames, for which tissue contrast was reduced across the B-scan. Mean A-line profiles also indicated that the tissue signals decreased with increased averaging, along with the reduction in the background (Fig. 5(f)).

### Computational refocusing improves resolution and deep tissue visibility

As the last step in the ultradeep OCT processing chain, computational refocusing with bandwidth control was applied to compensate for defocus away from the focal plane and to limit the spatial frequencies included in the reconstruction (Fig. 6). This combination allowed lateral resolution recovery while suppressing noise outside the system’s transverse spatial-frequency support, which matters most in weak-signal regions at depth.

**Figure 6.**
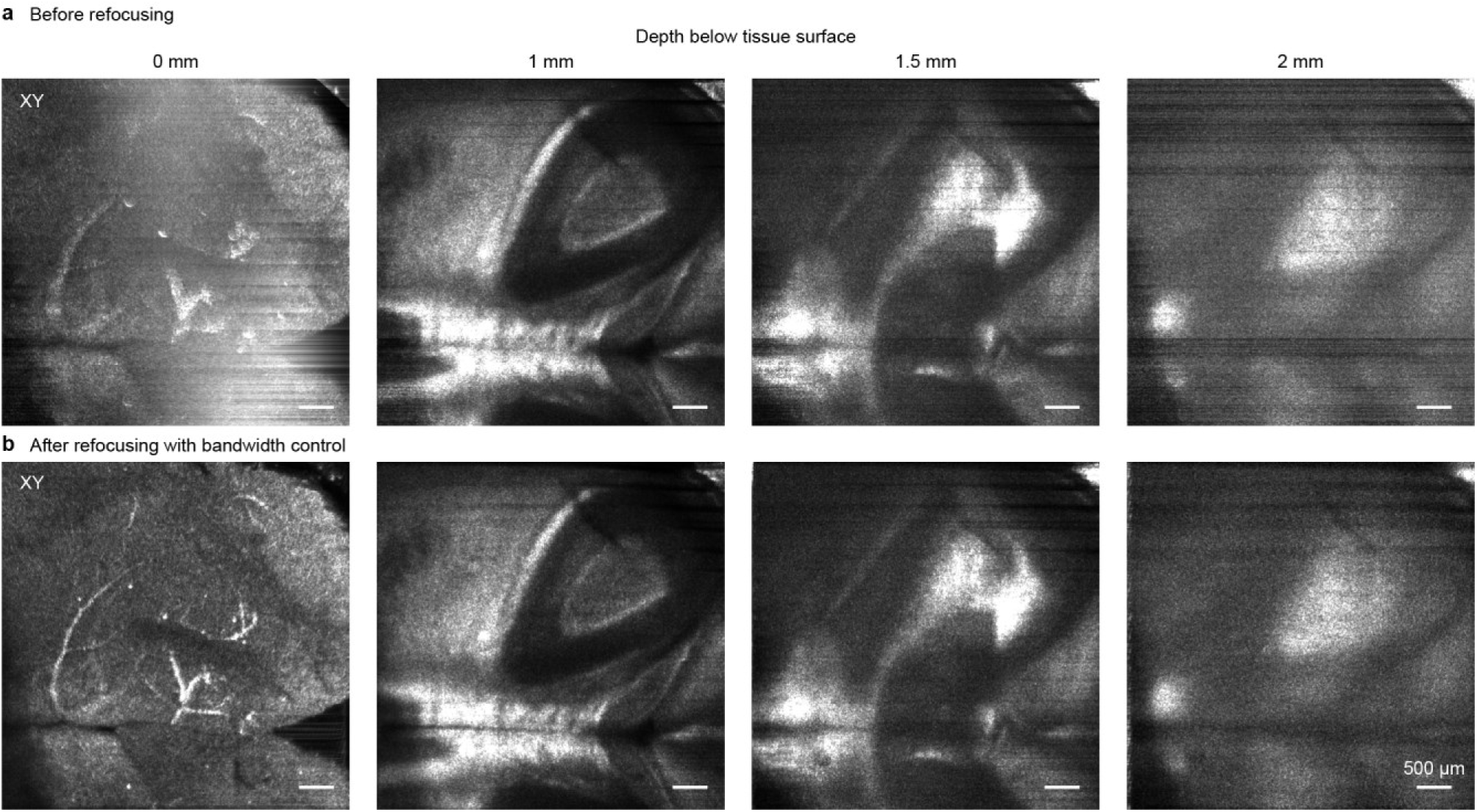
Computational refocusing with bandwidth control improves resolution and visibility of tissue features. Representative XY images in an *ex vivo* mouse brain, at depths of 0, 1, 1.5, and 2 mm below the tissue surface. **a,** Before refocusing. **b,** After refocusing with bandwidth control. Corresponding images at each depth are displayed with identical contrast settings. Scale bars, 500 µm.

The effect of refocusing, which improves both transverse resolution and SNR, was most apparent near the tissue surface (Fig. 6). Refocusing made the vasculature at the brain surface sharper and also enhanced structural details deeper within the tissue. At 1 mm depth, the refocused images showed sharper boundaries and improved definition of the curved and layered hippocampal structures. The improvements due to refocusing became less pronounced at depths of 1.5 and 2 mm. However, the bandwidth-control step along with refocusing modestly improved the localization and visibility of the remaining tissue features. Bandwidth control was particularly important at depth, where reduction of high-frequency noise improved visibility of low-contrast tissue structure. See S6 for additional details.

### Three-dimensional reconstruction of hippocampal subregions

The extended imaging depth achieved with ultradeep OCT made it possible to visualize the dorsal hippocampus in the acquired *ex vivo* mouse brain volumes (Figs. 1(c), 1(d) and 7). The hippocampus was bounded dorsally by the strongly scattering CC and ventrally by bright, myelin-rich regions of the thalamus. Owing to the higher OCT contrast of myelinated tissue, these surrounding structures provided anatomical landmarks for identifying the position and extent of the hippocampus.

Contrast within the hippocampus further facilitated segmentation of the CA and DG subregions. Segmentation and three- dimensional reconstruction in two representative mouse brains are shown in Fig. 7(a). The reconstructions exhibited the expected elongated, curved organization of the dorsal hippocampus. The CA extended along the dorsal aspect of the reconstructed volume, whereas the DG occupied the more ventral and internal regions. Insets showing the segmented structures within the corresponding OCT brain volumes depict the position and orientation of the segmented portion of the hippocampal formation within the surrounding tissue.

**Figure 7.**
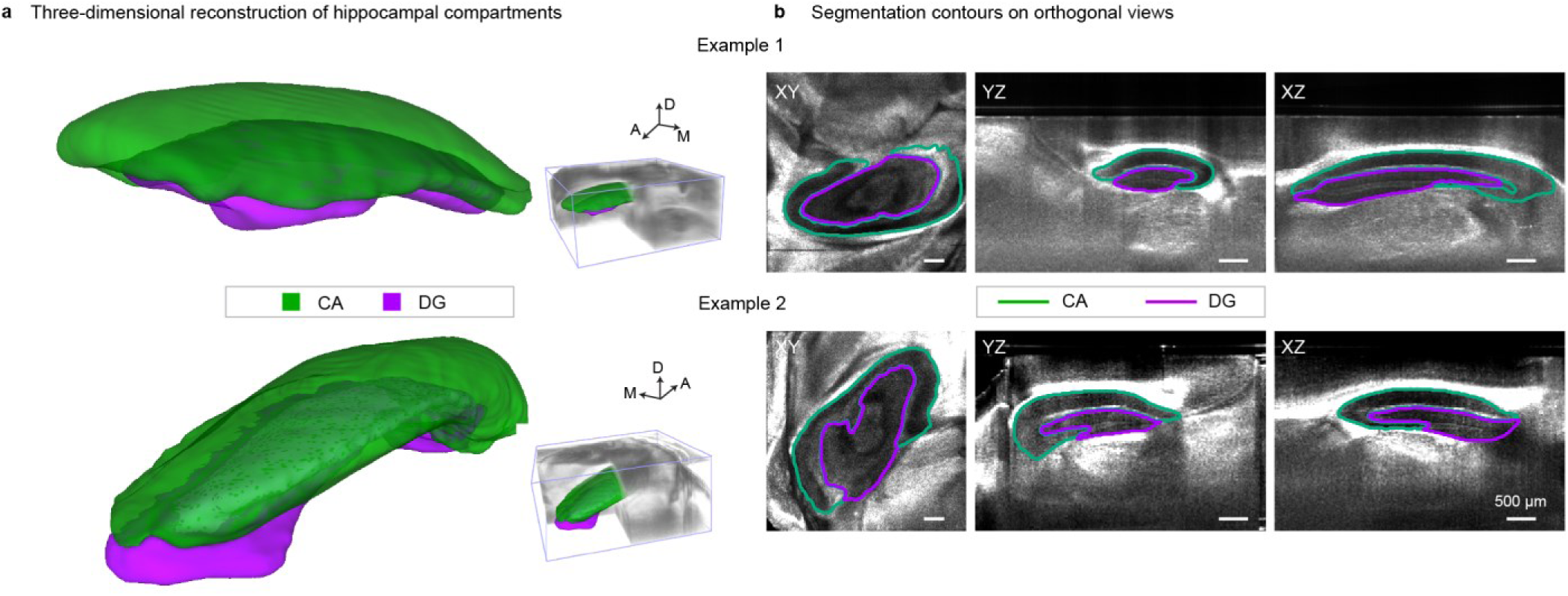
Three-dimensional segmentation and reconstruction of hippocampal subregions from ultradeep OCT volumes. **a,** Reconstructions from two representative mouse brain volumes, designated as Example 1 and Example 2. Segmented subregions are CA, green and DG, purple. The segmentations within the acquired OCT volumes (5 × 5 × 3.8 mm, X × Y × Z) are shown to the right. The 3D renderings are shown in perspective and carry no scale bar; see the orthogonal views in **b** for scale. A, anterior; D, dorsal; M, medial. **b,** Segmentation contours overlaid on representative orthogonal XY, YZ, and XZ OCT views. Scale bars, 500 µm.

Segmentation contours overlaid on orthogonal XY, YZ and XZ views illustrate the correspondence between the reconstructed boundaries and OCT tissue features (Fig. 7(b)). In the *en face* XY views, the contours followed the nested organization of the CA and DG within the hippocampus. The YZ and XZ views demonstrated that these boundaries could be followed through the depth and longitudinal extent of the dorsal hippocampus. See S7 for additional details.

### Imaging depths approaching 3 mm in chicken tissue

The complete ultradeep OCT imaging and processing pipeline was applied to chicken tissue to evaluate performance in a tissue type distinct from the mouse brain (Fig. 8). The XZ image showed detectable structural contrast throughout most of the acquired depth range. Corresponding *en face* images revealed heterogeneous tissue features at the surface and at depths of 1 and 2 mm. At the physical depth of 3 mm, the signal approached the background level, but residual spatially varying tissue contrast remained observable. The observed spatial variation and continuity at this depth were more consistent with tissue features than with AC artifacts or shadowing from superficial structures. Thus, ultradeep OCT provided a structural imaging depth approaching 3 mm in the representative chicken thigh result shown in Fig. 8. Across measurements in three chicken tissue samples, the structural imaging depth increased from 1.89 ± 0.09 mm for conventional OCT to 2.77 ± 0.26 mm for ultradeep OCT. Even greater imaging depths might be feasible as indicated by Fig. 5(b). See S5 for additional results in chicken tissue.

**Figure 8.**
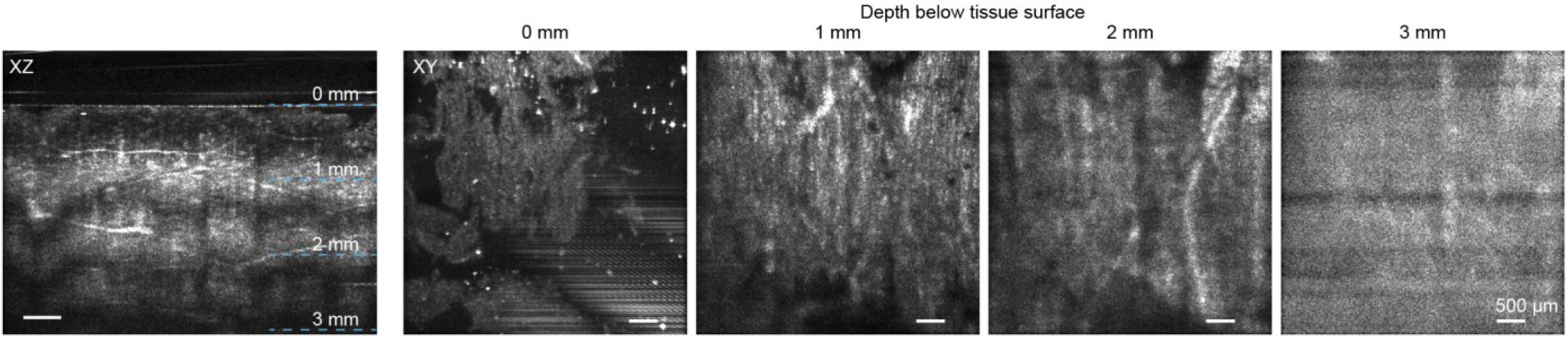
Ultradeep OCT imaging of chicken thigh tissue. Representative XZ cross-sectional OCT image and XY *en face* images acquired at depths of 0, 1, 2, and 3 mm below the tissue surface. Dashed blue lines in the XZ image indicate the locations of the XY images. Scale bars, 500 µm.

## Discussion

The imaging depths achieved with ultradeep OCT exceed those previously reported for structural imaging of fresh brain tissue, to the best of our knowledge [Roper, 1998; Bolmont, 2012; Srinivasan, 2012; Leahy, 2013; Choi, 2015; Chong, 2015; Chen, 2016; Yamanaka, 2016; Park, 2018; Yamanaka, 2019; Zhu, 2021]. Unlike in high-resolution imaging, where imaging depth is understood to be the depth at which diffraction-limited imaging is feasible, structural imaging depth here is understood to be the maximum physical depth at which tissue features remain detectable [Roper, 1998; Choi, 2015; Chong, 2015; Park, 2018]. Beyond the nominal structural imaging depths reported in this study, it was shown that ultradeep OCT volumes maintained sufficient contrast at depth to enable the identification and segmentation of hippocampal subregions. Ultradeep OCT could therefore enable the study of structural hippocampal changes in mouse models of neurodegenerative disease [Jullienne, 2022]. Deep cerebral angiography represents another potential application, including studies of the ischemic response [Chen, 2016].

However, the imaging depths reported here are not directly comparable to those reported by earlier studies. First, the depths reported here correspond to structural imaging rather than diffraction-limited, high-resolution imaging [Srinivasan, 2012; Zhu, 2021; Yoon, 2020]. Second, imaging depth is expected to differ between *in vivo* and *ex vivo* experiments. *In vivo* imaging depth depends significantly on the method of brain access. Imaging depth is expected to differ among open-skull access, glass cranial windows and preparations with and without removal of the dura [Roper, 1998; Srinivasan, 2012; Leahy, 2013; Choi, 2015]. Similarly, performance through a thinned-skull, reinforced window is expected to differ substantially from performance for transcranial imaging [Chong, 2015; Zhu, 2021; Wang, 2007; Kwon, 2023]. Additionally, the scattering coefficient of the rodent brain has been shown to increase with age [Chong, 2015] and imaging depth has also been shown to decrease postmortem [Choi, 2015]. Several previous *ex vivo* or *in vivo* studies used mice aged 5 to 12 weeks [Leahy, 2013; Choi, 2015; Chong, 2015; Park, 2018; Yamanaka, 2019], whereas the brains imaged in our study were extracted from mice that were 32 to 44 weeks old. On one hand, because brain scattering increases with age and has been reported to increase postmortem, ultradeep OCT could potentially achieve greater imaging depths *in vivo* in young mice than those reported in this study. On the other hand, the presence of dura, blood absorption, and physiological motion could all reduce the achievable imaging depth *in vivo*. Another consideration that makes the comparison of imaging depths imprecise is that although freshly extracted brains were imaged in this study, the exact acquisition time postmortem was not controlled for.

The coverslip placed on the brain during imaging provided a high-SNR signal for phase registration, while also approximating the optical configuration of an *in vivo* glass cranial window. However, the coverslip compressed the cortex, potentially altering deeper structures as well, and the results presented are for this distorted brain sample. Nevertheless, the demonstrated ability to image into the thalamus suggests that the ventral portions of the hippocampal formation may remain accessible even when the cortex is undistorted, as would be expected under *in vivo* conditions. This is another reason why depths reported here are not directly comparable to other studies. The maximum structural imaging depth demonstrated in chicken tissue also indicates that the imaging-depth improvement is reproducible across two types of biological tissues, although absolute imaging depth will likely remain dependent on tissue properties and structural composition.

Despite greater water absorption in the 1575 nm detection band than in the 1300 nm band, imaging depth was greater at 1575 nm in both mouse brain and chicken tissue (Fig. 2, SF5, and SF6). However, this is a system-level improvement, and differences in power, sensitivity, imaging range and resolution prevent a direct attribution to wavelength alone. Additionally, when the ultradeep processing steps were applied to the 1300 nm datasets, tissue features could be observed at the maximum depth that could be imaged. However, the contrast of tissue features at this depth was better in the 1575 nm result. Calculations indicate that, over most of the corresponding spectral range, the total attenuation coefficient of a 250 nm bandwidth centered at 1575 nm is lower than that of an equivalent bandwidth centered at 1300 nm, which supports the argument of greater imaging depth at 1575 nm. Other possible factors contributing to the greater imaging depth at 1575 nm include preferential attenuation of multiply scattered photons (owing to longer pathlengths) due to the increased water absorption [Kodach, 2010], and improved SNR at 1575 nm owing to its larger coherence volume [Harper, 2022]. See S5 for more details.

Greater imaging depths may be possible near 1700 nm, where attenuation is expected to be the lowest. However, practical system limitations must be considered. SD-OCT systems operating in this band are expected to have lower dynamic range because extended-range InGaAs sensors have higher noise levels than standard InGaAs sensors [Chong, 2015; Zhu, 2021; Thorlabs]. The quantum efficiency of standard InGaAs sensors decreases sharply past 1700 nm; therefore, the 1575 nm center wavelength could provide a more favorable balance than 1300 or 1700 nm for deep tissue imaging. The lower structural imaging depth reported for a 1700 nm swept-source OCT system [Park, 2018] than for a 1300 nm system [Choi, 2015] further emphasizes that imaging depth depends on overall system performance, including the dynamic range, rather than wavelength alone. See S5 for more details.

The benefit of reversing spectrometer roll-off for deep tissue imaging has been demonstrated previously [Spaide, 2008]. An additional advantage of the conjugate configuration is that it reduces the DR required to image a given depth range of the sample by selectively attenuating strong signals from shallow depths. The attenuation of superficial features, and the corresponding relative enhancement of deeper features, depends on the change in their positions between the standard and conjugate configurations. Thus, the conjugate imaging configuration effectively trades off SNR at shallow depths for increased SNR at larger depths. This approach is primarily relevant to SD-OCT because swept laser sources with negligible roll-off are available [Choi, 2015].

Balanced detection doubles the desired interferometric signal, while suppressing common-mode components of the Fourier domain OCT signal such as AC, source RIN, and any DC offset [Balakrishnan, 2026]. In the standard imaging configuration, the impact of AC artifacts can be mitigated by positioning the sample-air interface away from zero OPD. However, in the conjugate configuration, zero OPD is intentionally positioned deep within the sample, causing the AC artifacts to overlap with signals from deep tissue regions. AC artifacts were particularly prominent in this study because of the coverslip and the relatively low numerical aperture of the objective. Tilting the sample was therefore needed to reduce the intensity of the AC artifacts. However, significant AC artifacts persisted in brain samples even in the standard configuration; in the conventional OCT result in Fig. 1(c), the curved feature in the middle of the cortex marked by an arrow is an AC artifact. For clarity, the comparisons in Figs. 2 and 3 were presented after balanced detection; these images have less prominent AC artifacts and are therefore easier to interpret. The corresponding single-spectrometer results yield the same overall conclusions.

Balanced detection suppressed the AC artifacts but not completely, which made deep features in the conjugate configuration challenging to resolve. Balanced detection combined with AC subtraction was therefore critical for ultradeep imaging. AC frames acquired at a later time were ineffective in biological samples because subtle sample motion caused decorrelation. A shutter was used to acquire an AC-only frame immediately after the corresponding signal frames, but an AC-only frame could also be acquired in the middle of the acquisition sequence. A disadvantage of this AC subtraction approach was that an additional AC-only frame needed to be acquired at each Y position, which increased the total acquisition time.

Coherent averaging extended imaging depth by reducing the noise floor and is superior to incoherent averaging in this regard [Tomlins, 2007; Leartprapun, 2021]. The attenuation of tissue signals with prolonged averaging (Fig. 5(e), 5(f)) is likely caused by decorrelation of the signal due to motion in biological samples over time. Even in nominally stationary *ex vivo* tissue, sub-wavelength sample motions can alter the phase of the measured signal between frames, which cannot be corrected by the coverslip phase registration procedure. Under the imaging conditions used here, 16-frame averaging provided an effective compromise, lowering the noise floor and increasing the detectable depth while avoiding the signal loss observed with larger numbers of averaged frames. When the sample signal remains deterministic and noise varies randomly across repeated acquisitions, coherent averaging preserves the signal while suppressing random noise. It is therefore most effective at depths where the signal approaches or falls below the noise floor. Multiple scattering contributions are deterministic and are not suppressed by coherent averaging. In biological samples, decorrelation caused by motion could be mitigated by increasing the acquisition frame rate and compensating for sample motion. Coherent averaging is also expected to require correction of physiological motion when translating the ultradeep approach to *in vivo* imaging. However, shorter exposure times reduce the SNR of individual measurements, while reliable motion estimation is possible only at depths with sufficient SNR.

Access to the complex OCT signal allowed transverse resolution improvement by computational refocusing. In both brain and chicken samples, refocusing enhanced the sharpness of tissue features at a range of depths (Fig. 6 and SF5). This improvement was less evident at greater depths, likely because sample-induced aberrations and multiple scattering introduce wavefront errors that violate the assumptions underlying computational refocusing. Bandwidth control improved the visibility of deep features by filtering high-frequency noise. Although bandwidth reduction is theoretically expected to decrease resolution, no appreciable loss of resolution was observed here, likely because only very high spatial frequencies were filtered owing to the large relative bandwidth of the mask.

The SBR metric used to quantify imaging depth in *ex vivo* mouse brain volumes had several limitations. The SBR threshold was determined visually for each dataset. This approach was necessary because haze arising from multiple scattering and shadowing caused by bright superficial tissue structures often resembled genuine tissue features at depth. Consequently, subjective assessment based on structural continuity and texture was unavoidable. In addition, the bandwidth-control operation suppresses noise and thereby increases the SBR, which may make the final processing step in Fig. 1(b) appear to provide the largest relative improvement in imaging depth for this dataset. For these reasons, the SBR metric should be interpreted primarily as a tool for comparing imaging depth across processing techniques rather than as an absolute measure. The final imaging depths for ultradeep OCT and conventional OCT reported for each brain dataset were verified manually. The maximum imaging depth in chicken tissue was determined by visual inspection alone.

The reconstructions shown in Fig. 7 demonstrate segmentation feasibility but do not support a quantitative comparison between brains or genotypes. The volumes were manually segmented by a single observer using predefined anatomical landmarks; therefore, inter-observer variability was not assessed.

In conclusion, ultradeep OCT achieved sufficient imaging depth in fresh, uncleared *ex vivo* mouse brain tissue, enabling not only volumetric reconstruction of the dorsal hippocampus but also visualization of structures within the thalamus whose appearance was consistent with that of fiber tracts. With further development, ultradeep OCT could facilitate quantitative, longitudinal investigations of deep-brain structural changes in mouse models of neurodegenerative disease by enabling label-free visualization of hippocampal anatomy at a mesoscopic scale between MRI and optical microscopy. However, successful *in vivo* translation will likely face additional challenges that may require technical advances beyond those outlined in this study.

## Methods

### Balanced detection SD-OCT system with reference arm shutter

Light from a supercontinuum (SC) source (NKT SuperK Fianium FIU-6 OCT) was split using a broadband filter (NKT SuperK Split) and the spectral range 1150 to 2400 nm was coupled, after attenuation, into a fiber-based Mach-Zehnder interferometer. The interferometer consisted of a 90:10 coupler that split the light into reference (10%) and sample (90%) arms, two circulators that redirected the light returning from the sample and reference arms, and a 50:50 coupler that recombined the light from the two arms (Fig. SF1(a)). The sample arm contained a galvo scanner and a ∼0.1 NA, ∼40 mm working distance objective which was underfilled. The reference arm used optical elements similar to those in the sample arm, with the reference mirror placed after an optical shutter. The outputs of the 50:50 coupler were detected using two spectrometers with 250 nm spectral range and center wavelength of 1575 nm (Wasatch Photonics, Cobra 1600). The system axial resolution and imaging range in the brain tissue (refractive index, 1.35) were 10.1 µm and 3.78 mm, respectively. The transverse resolution was 14.5 µm. The incident power on the sample in the detected spectral band was approximately 25 mW, and the maximum sensitivity was measured to be 104.7 dB. The 1300 nm balanced detection system used for comparison also used the same supercontinuum source along with couplers and circulators designed for a center wavelength of 1300 nm and two 1300 nm spectrometers (Wasatch Photonics, Cobra 1300). See S1 for additional details.

### Mouse brain and chicken tissue samples

Brains from wild-type mice (n = 3, C57BL/6J, female, 32 to 44 weeks old) and one Alzheimer’s disease mouse (APP/PS1, male, 32 weeks old) were extracted and imaged immediately. All experimental animal procedures performed in this study were approved by the Institutional Animal Care and Use Committee at Cornell University. An FOV approximately centered over the posterolateral portion of a hemisphere was imaged (Fig. SF1(b)). Sections (n = 3) were obtained from chicken thigh or breast purchased from a local supermarket on the day of imaging. See S4 for more details.

### Acquisition and processing

A coverslip was placed on the tissue surface to serve as a phase reference surface [Ralston, 2007]. The sample, with the coverslip on top, was tilted to avoid spectrometer saturation and was imaged with the focal plane positioned deep in the tissue. For acquisitions in the conjugate imaging configuration [Spaide, 2008; Leartprapun, 2021], the zero OPD was positioned deep in the sample by adjusting the reference arm length until the upper coverslip surface was close to the maximum imaging range. Data from both spectrometers were acquired synchronously using a common frame trigger signal. Since the detected spectral ranges for the two spectrometers were not identical, the spectrometers were matched before calculating the balanced detection signal. The matching was performed using polynomial pixel resampling and amplitude matching curves determined using a calibration step that identified the optimal matching coefficients by minimizing the AC signal from a stack of coverslips [Balakrishnan, 2026]. The SC source was operated at 100% power level to obtain the lowest possible RIN. Additionally, an exposure time of 95 µs for the spectrometer camera, with a 10 kHz line rate, was chosen to further mitigate the impact of RIN. A BM-mode (repeated B-scans at one Y position) acquisition scheme was employed, where 16 frames were acquired at the same Y position for coherent averaging [Tomlins, 2007; Pfeiffer, 2020] followed by an AC-only frame, acquired with the shutter in the reference arm closed [Ai, 2006]. The scanned FOV was 5 × 5 mm, with 1024 A-scans per B-scan in X, 1024 B-scan positions along Y, and 17 repeated acquisitions at each Y position. The full volume acquisition took approximately 35 min. To assess coherent averaging performance (Fig. 5), a similar acquisition was made with chicken tissue, but without any Y position change. The coverslip results (Fig. 5(c)) were obtained similarly in the standard configuration.

Ultradeep processing consisted of balanced detection with AC subtraction, OCT reconstruction, coherent averaging, and computational refocusing with bandwidth control (Fig. SF2(a)). The first three steps were performed independently at each Y position, while refocusing was done after the entire coherently averaged volume was assembled. At every Y position, processing began by first subtracting, in the spectral domain, the AC-only frame from the signal frames for both spectrometers. Data from the second spectrometer were resampled and amplitude corrected using the previously obtained spectrometer calibration results. These matched spectra were then subtracted from the first spectrometer spectra to obtain the balanced signal. The balanced results were reconstructed by background subtraction, linear-in-*k* interpolation, dispersion compensation, and Fourier transform, after apodization with a Blackman window to obtain the complex OCT B-scans. For coherent averaging, the phase of all pixels in depth for each B-scan was registered to the phase of the upper coverslip surface to remove any system phase instabilities [Adie, 2012]. The coverslip phases across all B-scans were then registered, and coherent averaging was performed to produce a single, coherently averaged, complex B-scan at one Y position. The balanced detection with AC subtraction and coherent averaging steps were repeated at each Y position to obtain a 3D, coherently averaged, complex volume.

For computational refocusing, the coverslip phase was registered across all Y positions. After estimating the focal-plane position, *z_f_*, and the center of the transverse spectrum, computational refocusing was performed using the kernel: 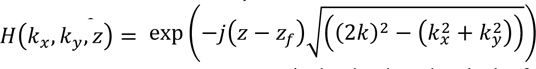, where *k* = 2*π*/*λ*, *λ* is the wavelength in the medium, *k_x_* and *k_y_* are the transverse wavevectors, *z* is the depth and *z_f_* is the focal plane position [Mulligan, 2019]. Bandwidth control was accomplished by apodizing the refocus kernel with a circular mask with Tukey-tapered boundary and a user-defined normalized radius, which was set to 0.9 for this study. The final result after balanced detection with AC subtraction, coherent averaging and computational refocusing with bandwidth control is termed the ultradeep OCT result. The coverslip was flattened on the final result to facilitate assessment of imaging depth and comparison among different datasets. Results obtained from scans made in the conjugate configuration were flipped vertically. The volumes were also resized axially to obtain isotropic voxels; each voxel was 4.883 µm on each side.

The volumes acquired in the mouse brain were intensity normalized in depth for improved visualization of the cross-sectional images using an approach inspired by depth-resolved attenuation coefficient estimation [Vermeer, 2013]. For pixels in a specified depth range, each pixel value was scaled by the inverse of the cumulative sum of amplitudes of all the pixels below it. This procedure provided a computationally efficient, A-line level depth normalization that produced an approximately uniform image contrast with depth, such that the entire *en face* stack could be viewed with a constant contrast setting.

The absolute pixel intensity values for images acquired with the 1575 nm and the 1300 nm systems were different. Similarly, intensity ranges for volumes acquired in the standard configuration were different from those acquired in the conjugate configuration. The refocusing with bandwidth control and the intensity normalization steps also modified the pixel intensity ranges. Contrast scaling used to display images is described in the figure captions. The contrast at each depth when showing XY images was set independently. See S1, S2, S3 and S6 for more details.

### Imaging depth assessment

Structural imaging depth was quantified by means of SBR profiles in depth on the OCT volumes without intensity normalization. SBR was calculated at each depth as the ratio of the mean amplitude in a signal region of interest (ROI) to the mean of the background amplitude in a separate background ROI, *SBR*(*z*) = 20 log_10_(*μ_signal_* (*z*)/*μ_bkgnd_* (*z*)). The signal and background ROIs were automatically determined on each *en face* plane as the brightest and darkest 50 × 50 pixel regions. ROIs were inspected and manually corrected, when necessary, to exclude imaging artifacts and to ensure that signal ROIs corresponded to locations containing tissue features. It was also confirmed that the background ROIs were placed in regions containing no tissue or, when this was not feasible, in regions containing noise only. Maximum structural imaging depth was defined as the depth at which SBR fell below a user-specified SBR threshold, which was determined for each dataset based on visual assessment of the final ultradeep OCT result. The same threshold was applied to determine the imaging depth for comparison among the different imaging steps. Dataset-specific thresholds ranged from 3.52 to 7.96 dB and were selected by visual assessment of structural continuity in the final ultradeep volume. This approach was found to reasonably match the visually determined imaging depth for each configuration. See S4 for more details.

### Segmentation of the hippocampus

Segmentations of the dorsal hippocampus in two *ex vivo* mouse brain volumes were performed manually in 3D Slicer [Pinter, 2019]. The hippocampus was segmented first, based on the boundaries of the surrounding myelin-rich structures. Next, the SLM within the hippocampus, which appeared as a relatively bright band, was identified. The hippocampus was segmented into its two principal subregions: the CA, comprising strata oriens, pyramidale, radiatum and lacunosum-moleculare, and the DG, comprising the molecular and granule cell layers and the hilus. The CA-DG boundary was placed along the ventral edge of the bright SLM band. Laterally, near the CA3-hilus junction, where the SLM contrast was insufficient, the boundary was placed by geometry. The subfields CA1, CA2 and CA3 were not individually resolved. The segmentation outlines on the three orthogonal views were used to assess the spatial continuity and anatomical consistency of the segmentations throughout each OCT volume. Finer laminar features, as well as the hilus, were also visible but were less consistent. Example 1 in Fig. 7 is from an Alzheimer’s model mouse and example 2 is from a wild-type mouse brain. See S7 for more details.

## Data availability

The data supporting the findings of this study are available from the corresponding author upon reasonable request.

## Code availability

Custom code used in this study is available from the corresponding author upon reasonable request.

## Acknowledgements

We gratefully acknowledge funding support from the National Institutes of Health (R01EB031226). We sincerely thank Kristine E. Kolkman and Niranjan V. Kulkarni for providing the excised mouse brain samples for this study.

## Author contributions

S.G.A. initiated and supervised the project. S.B. and G.A. designed and built the experimental setup. G.A. performed initial feasibility experiments that informed the final experimental design and identified limitations addressed in the final acquisition approach. S.B. designed the final acquisition scheme, performed the imaging experiments reported in the manuscript, developed the post-processing tools, and analyzed the experimental results. S.B. prepared the original draft. S.B. and S.G.A. discussed the results and finalized the manuscript. All authors approved the submitted version.

## Supplementary material

### S1: 1575 nm and 1300 nm balanced detection SD-OCT systems

The balanced detection SD-OCT systems at the two center wavelengths of 1300 nm and 1575 nm (Fig. SF1(a)) utilized a fiber-based Mach-Zehnder interferometer (MZI). The interferometer components (fiber couplers and circulators) and the spectrometers were different for the systems at 1300 nm and 1575 nm. The components in the sample and reference arms were identical in the two systems. The NKT SuperK Fianium FIU-6 OCT supercontinuum (SC) source had an approximate output spectral range from 400 to 2400 nm. A spectral splitter (NKT SuperK Split) was used to divide the output spectrum of the SC into a visible wavelength band from 400 to 1150 nm and a near-infrared (NIR) band with wavelengths from 1150 to 2400 nm. The NIR output of the splitter was attenuated using a neutral density filter and coupled into the MZI using a fiber collimator (NKT SuperK Connect). The supercontinuum source was operated at 100% power level to obtain an optimal spectrum and the lowest relative intensity noise (RIN) possible. The spectrum coupled into the interferometer extended from 1150 to 2400 nm, while the detected spectral ranges were 1450 to 1700 nm for the 1575 nm system and 1170 to 1420 nm for the 1300 nm system. The axial resolution and imaging range for the 1300 nm system in brain tissue (RI = 1.35) were 6.8 µm and 2.55 mm, respectively. The lateral resolution for the 1300 nm system was 11.8 µm. The maximum sensitivity for the 1300 nm system was 98.6 dB. The 6 dB roll-off distance, in tissue, for the 1300 nm system was 1.36 mm and for the 1575 nm system was 1.46 mm. The roll-off distance was determined by fitting a second order polynomial to a translated-mirror dataset. The same model was used to plot the roll-off curve in Fig. 3(a). The measured dynamic range for the 1300 nm system was 63.7 dB and for the 1575 nm system was 61.9 dB. The variable attenuator before spectrometer A was used to coarsely match the detected signal levels on the two spectrometers.

The total power incident on the sample in the 1150 to 2400 nm range, as measured by a thermal power sensor, was approximately 44 mW. To estimate the power incident on the sample in the detected spectral ranges, the incident power was measured with narrowband filters centered at 1300 nm or 1550 nm and the thermal power sensor. This measurement was then scaled based on detected spectra for each system. The incident power in the detection range was estimated to be 25 mW for the 1575 nm setup and 9 mW for the 1300 nm system. The system was designed with a broad incident spectrum to enable direct comparisons at 1300 and 1575 nm. A disadvantage of this scheme was that a significant portion of the optical power incident on the sample fell outside the detected spectral range and therefore did not contribute to the OCT signal. In future implementations, only the spectral band used for detection, approximately 1450 to 1700 nm, will be coupled into the interferometer and incident on the sample.

**Figure SF1.**
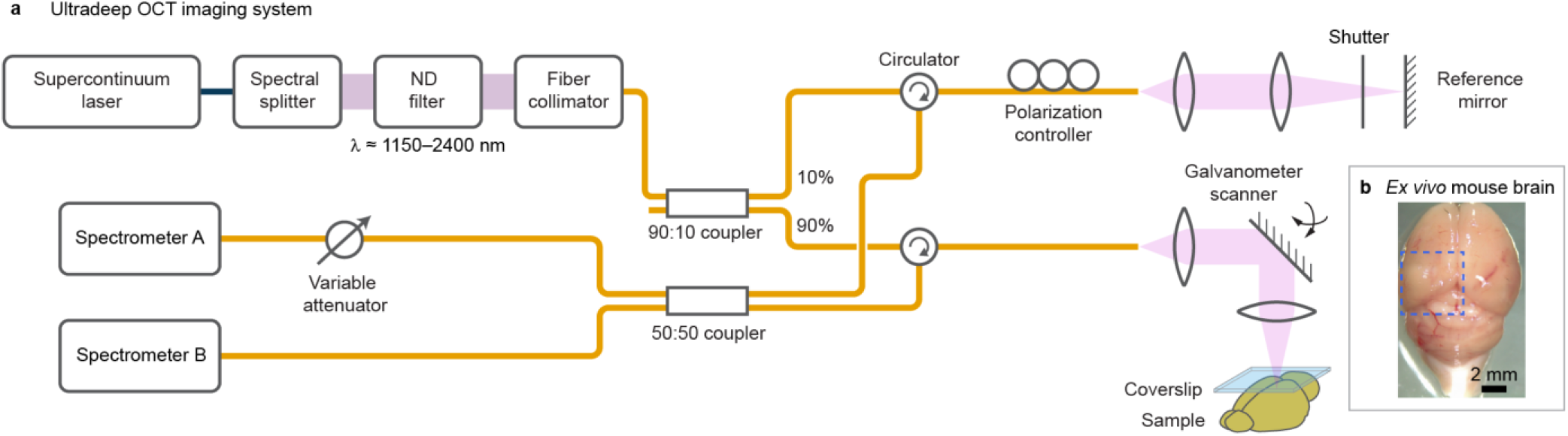
Ultradeep OCT imaging system. **a,** Schematic of the ultradeep OCT system. Broadband light from a supercontinuum laser is spectrally filtered to approximately 1150 to 2400 nm and coupled to a fiber-based Mach-Zehnder interferometer configured for balanced detection. The fiber couplers, circulators and spectrometers had wavelength ranges appropriate for operation at a center wavelength of either 1300 or 1575 nm. **b,** Photograph of an *ex vivo* mouse brain, showing a representative 5 × 5 mm scanned FOV outlined by the blue dashed box. Scale bar, 2 mm.

For BM-mode acquisition, 16 repeated B-scans were acquired at the same Y location with the reference-arm shutter open, followed by an AC-only frame acquired with the shutter closed. This sequence was repeated at the successive positions, until the specified number of Y positions had been acquired. Custom LabVIEW software was used to synchronize galvanometer scanning, shutter actuation and spectrometer data acquisition. A common external trigger was used to start data acquisition from both spectrometers.

For a typical experiment, the sample was transferred to a dish, and a coverslip (#1.5, 170 µm thickness) was gently placed on the sample surface. The sample was positioned such that the coverslip was near the top of the imaging range, and the focal plane was approximately in the middle of the imaging range. In the case of brain samples, these steps resulted in a focal plane positioned below the corpus callosum (CC). The sample tilt was adjusted to prevent spectrometer saturation. The reference arm length was then adjusted to image the sample in the conjugate configuration. Finally, the sample tilt and the reference power were fine-tuned before the data acquisition. BM-mode acquisition in the conjugate configuration at 1575 nm was performed for all samples. Data from this acquisition was processed to produce the single-spectrometer detection result, the balanced detection with AC subtraction result, the coherent averaging result and the final ultradeep OCT result after computational refocusing with bandwidth control. Regular volume acquisitions, with only one frame per Y position and no AC-frame, in the standard configuration at 1575 nm, as well as BM-mode and regular acquisitions at 1300 nm were primarily used to verify the improved performance of the ultradeep approach and were therefore not repeated each time. For comparisons among configurations, the scanned FOV and the focal plane position were kept unchanged. The approximate scanned FOV position on an *ex vivo* mouse brain sample is shown in Fig. SF1(b).

### S2: Ultradeep OCT processing

The steps involved in reconstructing ultradeep OCT results are summarized in Fig. SF2(a). The processing pipeline can be split into four groups of steps, which are identified by different colors. The steps involved in producing the balanced detection signal are shown in blue. At each Y position, the first step involved subtracting the AC frame from the BM-mode frame in the spectral domain. Spectrometer matching calibration performed separately, as described previously [Balakrishnan, 2026], generated a pixel resampling vector and an amplitude matching curve. The data from spectrometer B was resampled and amplitude corrected using these calibration results and was then subtracted from the spectrometer A data to produce the balanced detection with AC subtraction signal. Background was then subtracted, using the median of the A-scans as the background estimate. The resulting spectral domain signal was resampled to be linear in wavenumber (*k*), followed by numerical compensation of the dispersion mismatch between the sample and reference arms. The spectrum was then apodized using a Blackman window and Fourier transformed to generate an OCT B-scan. These standard OCT reconstruction steps, depicted by green boxes, generated the complex, balanced detection with AC subtraction B-scan. To produce the single-spectrometer result, the same steps were repeated, except for the AC subtraction and balanced detection step, using data only from spectrometer A.

**Figure SF2.**
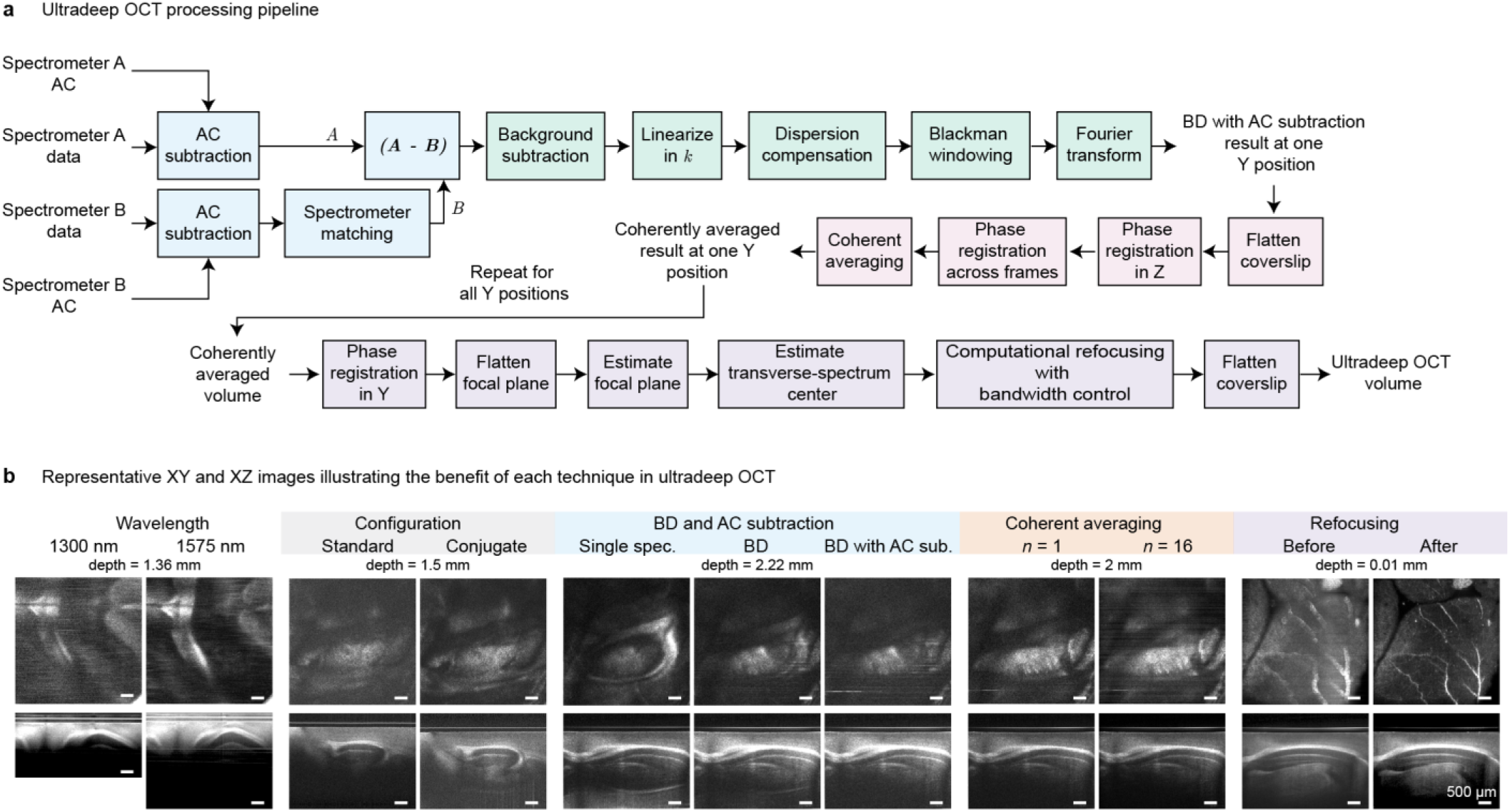
Ultradeep OCT processing pipeline and representative intermediate results. **a,** Processing pipeline for ultradeep OCT. Steps involved in balanced detection are in blue, OCT reconstruction steps are colored green, operations required for coherent averaging are in pink and computational refocusing with bandwidth control processing steps are in purple. **b,** Paired *en face* (top row) and cross-sectional (bottom row) images corresponding to the principal acquisition and processing comparisons. Contrast was adjusted independently for each image in the wavelength and configuration groups and was identical within all other groups. Wavelength group images are for the WT2 dataset. All other images are for the AD dataset. BD, balanced detection; AC, autocorrelation; *n*, number of averaged frames. Scale bars, 500 µm.

For coherent averaging of the repeated acquisitions at one Y position (pink boxes), the tilted coverslip surface was first made flat, followed by phase registration in the axial direction using the upper coverslip surface as the phase reference [Adie, 2012]. Next, the coverslip phase in each repeated B-scan was registered to the phase of the first frame. These registered, complex B-scans were then coherently averaged to give the result at the current Y position. These steps were repeated at every Y position, and the resulting B-scans were combined to produce the coherently averaged three-dimensional volume.

Computational refocusing (purple boxes) began with registration of the coverslip phase along the Y direction. The coverslip flattening process, performed as a part of coherent averaging, caused the focal plane to become tilted. Therefore, the flattening of the coverslip was first undone to make the focal plane flat again. The axial position of the focal plane was estimated from the minimum of the root-mean-square radius (RMSR) of the transverse spatial-frequency spectrum. The RMSR profile as a function of depth was calculated as 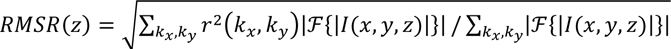, where 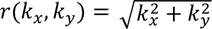 and ℱ{⋅} denotes the Fourier transform operator. The focal plane position, *z_f_*, was estimated as the location of the local minimum in the smoothed RMSR profile within the sample region, as shown in Fig. SF8(a) and SF8(b). The center of the transverse spatial-frequency spectrum was then estimated to correctly position the defocus-correction kernel (Fig. SF8(c)). Computational refocusing was performed using a kernel similar to the angular-spectrum propagation kernel [Adie, 2012; Mulligan, 2019]. Bandwidth control was implemented by pre-multiplying the refocus kernel by a circular mask with a Tukey-tapered boundary (Fig. SF8(d)). Finally, the coverslip surface was flattened again to facilitate visualization, producing the final ultradeep OCT volume. The reconstructed volumes were resized in Z to obtain isotropic voxels with a side length of 4.883 µm, calculated as 5000/1024 µm. Acquisitions made in the conjugate configuration were also flipped vertically for display.

The ultradeep OCT brain volumes were intensity normalized in depth using a method similar to that introduced by Vermeer et al. [Vermeer, 2013] for depth-resolved estimation of attenuation coefficients. For pixels in the depth range *r* ∈ [*r_min_*, *r_max_*], the pixel value at depth *r* and transverse coordinates (*p*, *q*) was replaced by *I*^′^[*p*, *q*, *r*] = *I*[*p*, *q*, *r*]/*f*. The denominator *f* was a regularized, normalized, cumulative sum of pixels below *r*: 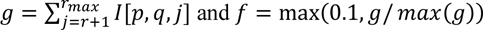 and *f* = max(0.1, *g*/ *max*(*g*)). The pixel values of the intensity-normalized volume were finally rescaled to match the input volume pixel value range. The shallower depth limit (*r_min_*) prevented excessive attenuation of structures near the tissue surface, whereas the deeper depth limit (*r_max_*) prevented progressive amplification of the noise at large depths. This procedure provided computationally efficient, A-line level depth normalization with only two user-selected depth parameters. Based on comparisons of the mean intensity profiles and visual image quality, this method produced more uniform contrast with depth than gamma compression or percentile-based intensity scaling.

Fig. SF2(b) illustrates each step of the ultradeep OCT approach using paired XY and XZ images in which one imaging or processing condition was changed at a time. The wavelength group of images compares results in the standard imaging configuration at 1300 and 1575 nm. The XZ images have different Z sizes because the imaging range of the 1575 nm system was larger. The XY images at the same depth in the sample for both wavelengths showed better feature contrast in the 1575 nm case. The 1575 nm XZ image also showed features at depths beyond the imaging range of the 1300 nm system. The next comparison illustrates that changing from the standard to the conjugate imaging configuration at 1575 nm increased the detected signal at depth. However, the combination of conjugate imaging configuration, coverslip and bright myelinated regions in the brain produced strong AC artifacts that overlapped with tissue features at depth. Balanced detection with AC subtraction significantly suppressed these artifacts and enhanced visibility of low SNR features at depth. The coherent averaging comparison shows that averaging 16 registered frames reduced the noise floor and improved the contrast for deeper structures relative to no averaging. The final group demonstrates the benefits of computational refocusing with bandwidth control. This step increased the transverse resolution of brain features close to the surface, as seen on the XY images. The bandwidth-control operation is effectively a low-pass filter in the spatial frequency domain and improved SNR by suppressing high-frequency noise. Display settings were kept the same for each type of image in a group, except for the wavelength and the configuration groups, where the contrast on each image was set independently. The impact of depth-dependent intensity normalization is also illustrated with the cross-sectional images. The images in the wavelength group are displayed without intensity normalization. Those in the configuration, balanced detection and coherent averaging groups are also displayed with no intensity normalization, but with *γ* = 0.5. The cross-sectional images in the refocusing group are displayed with depth-dependent intensity normalization. The contrast on these images is more uniform, allowing better visualization of features in depth.

### S3: Balanced detection with AC subtraction

The benefits of combined balanced detection and AC subtraction are illustrated in Fig. SF3. In a coverslip stack sample, prominent AC artifacts that were observed in the single-spectrometer result (magenta arrow, Fig. SF3(a)) were strongly attenuated by balanced detection, although weak residual artifacts remained (orange arrow, Fig. SF3(a)). These residual artifacts were removed when balanced detection with AC subtraction was used. The mean A-line plot, shown on the bottom row, illustrates that the AC artifact peak level near z = 70 was 87 dB in the single-spectrometer case and 77 dB for balanced detection alone. For balanced detection with AC subtraction, this peak was suppressed to 56 dB, which was the level of the noise floor. Additionally, the sample-associated signal peak values near z = 430 were 90, 95 and 95 dB while the noise floors were approximately 54, 55 and 56 dB for the three conditions, denoting a 3 dB improvement in SNR when performing balanced detection with AC subtraction over the single-spectrometer case.

A similar trend was observed in the mouse brain B-scans shown in Fig. SF3(b). Balanced detection markedly attenuated the AC artifacts from the coverslip and the CC, but weak residual artifacts remained visible at depths close to the zero OPD. Balanced detection with AC subtraction suppressed the residual artifacts. The mean A-line profiles show that the AC artifact peak near z = 50 was reduced from 80 to 55 dB and the signal peak near z = 480 increased from 60 to 62 dB when comparing the balanced detection with AC subtraction result with the single-spectrometer result.

**Figure SF3.**
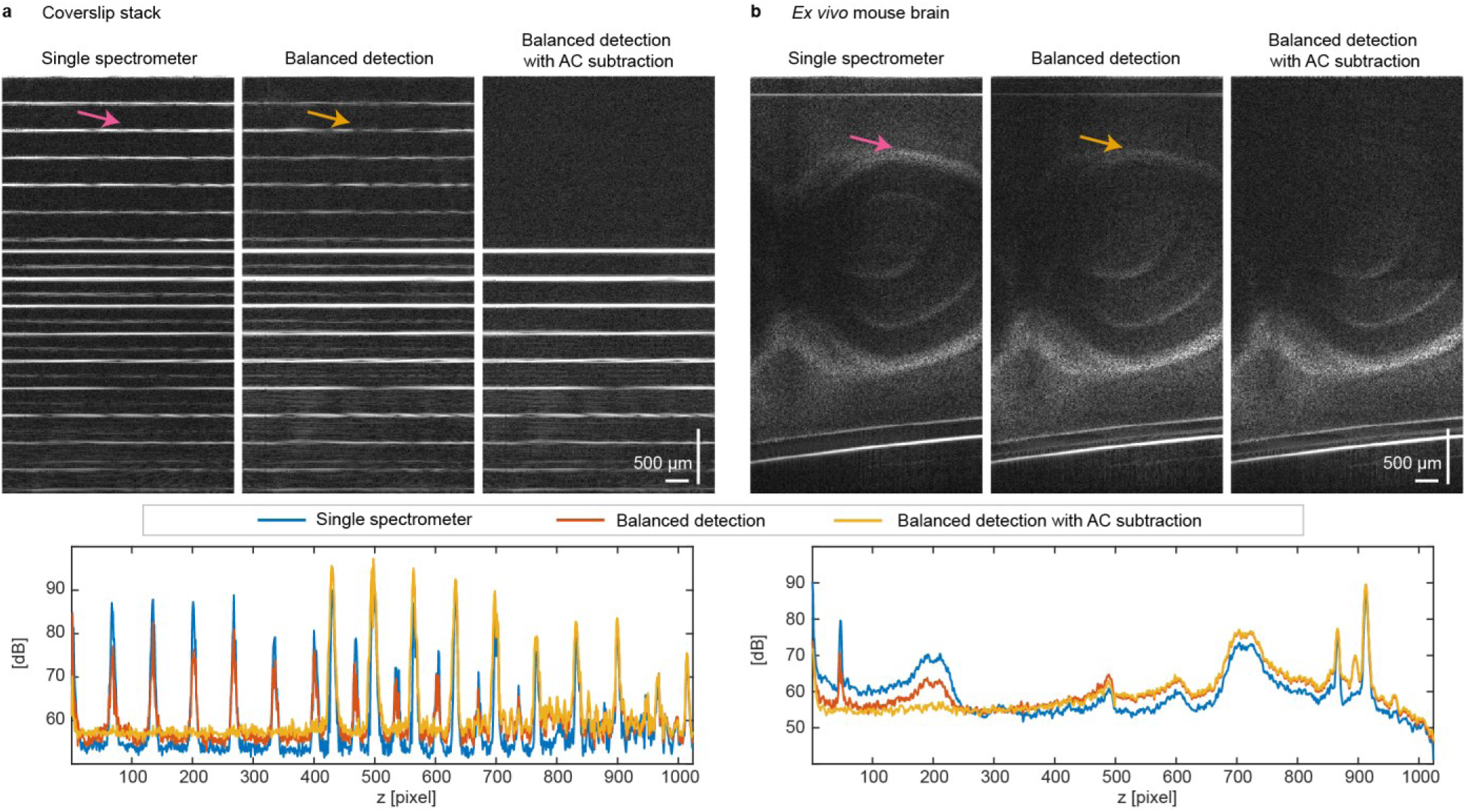
AC artifact suppression and signal peak improvement by balanced detection with AC subtraction. **a,** Measurements of a coverslip stack, acquired in the standard configuration. **b,** B-scans of an *ex vivo* mouse brain, acquired in the conjugate configuration, shown without vertical flip or coverslip flattening. **Top**, Representative B-scans obtained using a single spectrometer, balanced detection alone, and balanced detection combined with AC subtraction. Images within each group are shown without depth-dependent intensity normalization, with identical contrast settings and *γ* = 0.5. Magenta arrows identify AC artifacts on the single-spectrometer B-scans; orange arrows mark the residual artifacts after balanced detection. **Bottom**, Corresponding mean A-line profiles calculated by averaging 64 A-lines.

### S4: Assessment of imaging depth

Structural imaging depth was quantified by an SBR metric. The SBR metric as a function of depth was calculated using mean signal and background amplitudes measured in separate ROIs on the *en face* image at each depth. The signal and background ROIs were automatically determined as the brightest and darkest 50 × 50 pixel regions on the image (Fig. SF4(c)). The ROIs were inspected and manually corrected when necessary to ensure that imaging artifacts were excluded and that the signal ROI contained tissue features. The background ROIs were checked to ensure that these corresponded to regions without any tissue or to regions deep enough in tissue where there was only noise. SBR was defined as the ratio of the mean amplitude of the signal ROI to the mean amplitude in the background ROI, *SBR*(*z*) = 20 log_10_(*μ_signal_* (*z*)/*μ_bkgnd_* (*z*)).

SBR profiles for volumes with different techniques compared against ultradeep processing are shown in Fig. SF4(b). Maximum structural imaging depth was defined as the depth at which SBR fell below a dataset-specific threshold. This SBR threshold was determined for each dataset based on visual assessment of tissue feature visibility on *en face* images of the final ultradeep OCT result. The same threshold was applied to all imaging configurations for that dataset. This approach is relatively insensitive to differences in absolute signal and background levels and, among the evaluated approaches, provided the closest agreement with manual assessments of imaging depth for each configuration, while requiring only one user-defined parameter.

Comparison of SBR-limited imaging depths for each technique in ultradeep OCT was performed on datasets acquired with the same FOV. Signal and background ROIs were first determined from the final ultradeep OCT volume. These ROIs were then used to calculate the SBR at each depth for the coherently averaged, balanced detection with AC subtraction, and single-spectrometer detection volumes. Signal and background ROIs on the scans with the standard imaging configuration, at both 1575 nm and 1300 nm, were determined separately for SBR calculation as the sample positioning was different in these cases. SBR profiles for all configurations were plotted as a function of physical depth, with the tissue surface defined as 0 mm. For each configuration, the depth at which its SBR curve fell below the SBR threshold was taken as the maximum structural imaging depth. The final imaging depths for ultradeep OCT and conventional OCT reported for each brain dataset were verified manually. The maximum imaging depth in chicken tissue was determined by visual inspection alone as it was not possible to reliably identify background ROIs at each depth.

Ultradeep imaging was performed on four *ex vivo* mouse brains, designated AD, WT1, WT2, and WT3. Both hemispheres of WT2 were imaged, resulting in five datasets overall. Fig. SF4(a) summarizes the maximum structural imaging depth for each technique in the ultradeep OCT approach. The summary statistics were calculated from two measurements for conventional OCT at 1300 nm, from four measurements for OCT at 1575 nm in the standard configuration and from five measurements for the remaining techniques.

The SBR profiles as a function of depth for the different configurations in WT1 are shown in Fig. SF4(b). The SBR threshold in this case was 7.04 dB (ratio of 2.25), resulting in a maximum imaging depth of 1.4 mm for conventional OCT and 2.5 mm for ultradeep OCT. The increase in SBR at depths greater than 2.5 mm for 1575 nm, conjugate configuration (green curve) was caused by uncorrected AC artifacts. These artifacts were suppressed by the balanced detection with AC subtraction step (pink curve). The SBR thresholds used for the other volumes were 3.52, 3.52, 3.52, and 7.96 dB (ratios 1.5 and 2.5).

**Figure SF4.**
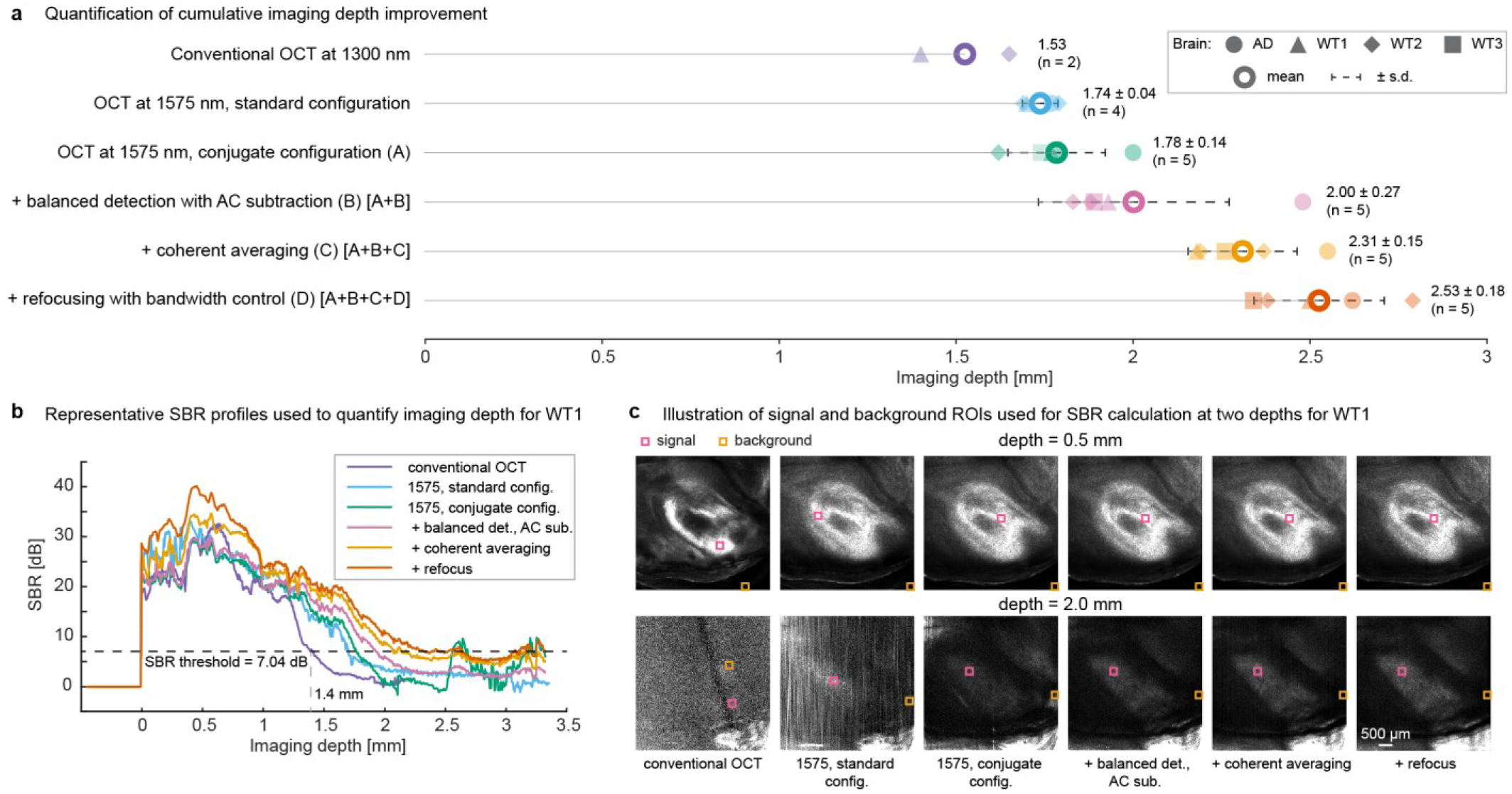
Structural imaging depth assessment. **a,** Imaging depth after implementation of the techniques used in ultradeep OCT. Measurements were obtained from five volumes acquired in four *ex vivo* mouse brains; both hemispheres were imaged for WT2. Filled symbols identify individual brain samples, and large open circles with error bars indicate mean ± s.d. Available sample sizes are shown for each technique. **b,** SBR-versus-depth profiles for WT1. The horizontal dashed line indicates the SBR threshold. The increase in SBR after 2.5 mm in the 1575 nm, conjugate configuration profile was caused by AC artifacts. **c,** Signal and background ROIs used for SBR calculation at depths of 0.5 and 2.0 mm. Pink and orange boxes denote signal and background ROIs, respectively. Scale bar, 500 µm.

### S5: Better penetration depth with the 1575 nm system

Although the improvement in imaging depth in OCT at 1300 nm relative to 800 nm is well established [Roper, 1998; Carrion, 2007; Marchand, 2018], investigations at wavelengths longer than 1300 nm have focused primarily on 1700 nm [Chong, 2015; Yamanaka, 2016; Park, 2018; Yamanaka, 2019; Zhu, 2021]. Several studies have nevertheless reported comparable or improved imaging performance near 1575 nm relative to 1300 nm, including in biological samples with high water content [Bouma, 1998;

Carrion, 2007; Biedermann, 2009; Kodach, 2010; Xu, 2014]. Despite these findings, the spectral region near 1575 nm has received comparatively limited attention, largely because of the strong water absorption peak near 1450 nm (Fig. SF7(b)).

In this study, the imaging depth was found to be greater with the 1575 nm system as compared to the 1300 nm system in both the *ex vivo* mouse brain and chicken tissue samples. Fig. 2 demonstrates better results at 1575 nm by comparing balanced detection results at the two wavelengths. Comparisons of regular acquisition standard configuration volumes, as well as volumes with the entire ultradeep OCT processing at 1300 nm versus 1575 nm were also performed. The maximum structural imaging depth was found to be greater at 1575 nm. Representative results in chicken tissue are shown in Fig. SF5. Compared with conventional 1300 nm OCT, the complete ultradeep OCT approach depicted the tissue features in the XY frame at a depth of 1.84 mm more clearly. The dark and bright regions visible on the 1300 nm *en face* image are not true tissue features but are caused by shadowing from bright superficial structures.

**Figure SF5.**
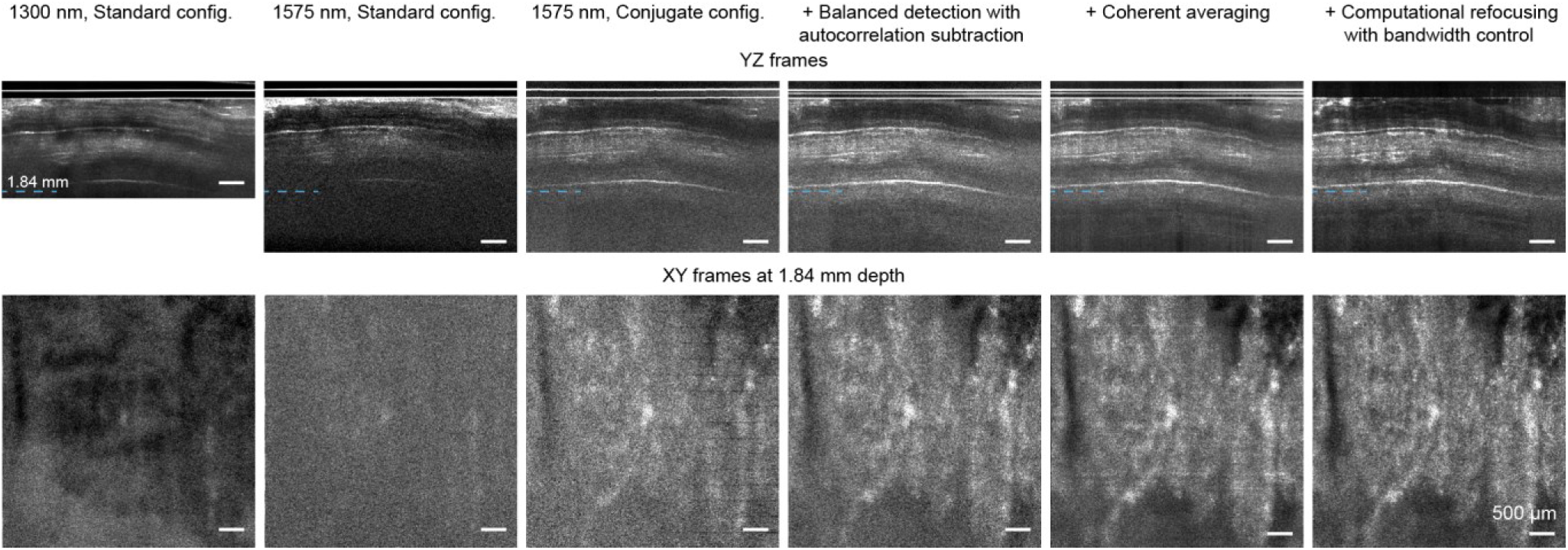
Ultradeep OCT results in chicken tissue. Representative cross-sectional frames (top row) and corresponding XY *en face* frames acquired at a depth of 1.84 mm (bottom row) illustrate results with the different techniques used in ultradeep OCT. From left to right: conventional OCT, 1575 nm OCT with the standard configuration, 1575 nm OCT with the conjugate configuration, balanced detection with autocorrelation subtraction, coherent averaging, and computational refocusing with bandwidth control. The blue dashed lines in the cross-sectional frames indicate the depth at which the *en face* images are shown. Contrast on each image was set independently. Scale bars, 500 µm.

The effect of longer center wavelength was clearer when comparisons of the complete ultradeep OCT acquisition and processing pipeline at both 1300 nm and 1575 nm were made (Fig. SF6). Data were acquired with the two systems in the conjugate configuration and processed using balanced detection with AC subtraction, 16-frame coherent averaging, and computational refocusing with bandwidth control. In the *ex vivo* mouse brain, the XY *en face* frame at a depth of 1.98 mm showed clearer visualization of the tissue region below the hippocampus at 1575 nm than at 1300 nm (Fig. SF6(a)). In chicken tissue, structural features remained visible near the maximum imaging range of the 1300 nm system (Fig. SF6(b)). However, comparison of the XY frames at the same depth of 1.84 mm showed that tissue features, indicated by arrows on the XY images, are better resolved in the 1575 nm result. The bright and dark patches in the 1300 nm chicken-tissue *en face* image, which give the impression of better tissue contrast, are once again caused by shadowing associated with superficial tissue structures.

**Figure SF6.**
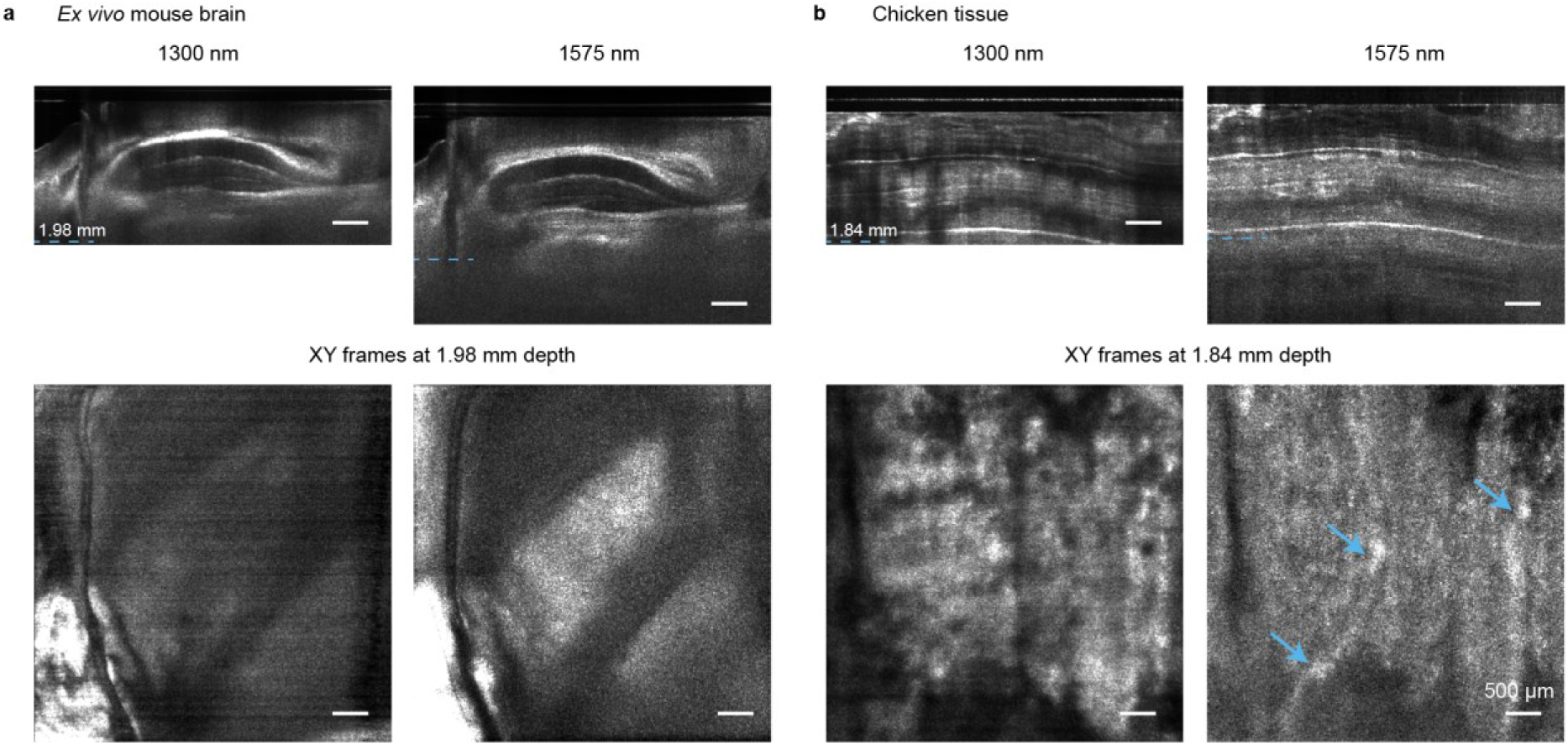
Improved visualization at depth with the 1575 nm system compared with the 1300 nm system when using the ultradeep OCT acquisition and processing pipeline. **a,** Representative cross-sectional frames (top row) and corresponding XY *en face* frames (bottom row) from *ex vivo* mouse brain acquired with the 1300 nm and 1575 nm systems. **b,** Corresponding comparison in chicken tissue, with XY frames at 1.84 mm depth. For both wavelengths, data were acquired in the conjugate configuration and processed using balanced detection with AC subtraction, 16-frame coherent averaging, and computational refocusing with bandwidth control. Blue dashed lines in the cross-sectional frames indicate the depths at which the corresponding XY frames were extracted. Blue arrows on XY frame in **b** indicate tissue features that are better seen on the 1575 nm image compared with the XY image at 1300 nm. Contrast on each image was set independently. Scale bars, 500 µm.

The calculated wavelength-dependent scattering (*μ_s_*), absorption (*μ_a_*) and total attenuation (*μ_t_*) coefficients for brain tissue are shown in Fig. SF7. The tissue model assumed water and lipid volume fractions of 75% and 15%, respectively. Scattering coefficients were obtained from the values in Table 2 of Ref [Jacques, 2013], assuming the scattering anisotropy (*g*) to be 0.85. Water absorption coefficients were obtained from Ref [Kou, 1993] and lipid absorption coefficients were obtained from Ref [Anderson, 2006]. The total attenuation coefficient was calculated as, *μ_t_* = *μ_s_*_,*brain*_ + *p_water_μ_a_*_,*water*_ + *p_lipid_μ_a_*_,*lipid*_. Fig. SF7(a) compares *μ_t_* over 250 nm wide wavelength ranges centered at 1300 nm and 1575 nm, corresponding to the spectral detection range of the spectrometers used in this study. The water absorption peak at 1450 nm increases the total attenuation near the short wavelength edge of the 1575 nm detection band and also near the long wavelength edge of the 1300 nm detection range. Nevertheless, *μ_t_* in the 1575 nm case is lower than that in the 1300 nm band over much of the corresponding range. The average values of *μ_t_* are approximately 4.6 mm^−1^ and 4.9 mm^−1^ in the 1575 nm and 1300 nm bands, respectively.

Fig. SF7(b) shows *μ_t_*, *μ_s_* and *μ_a_* in the wavelength range from 800 to 2000 nm, together with the 250 nm wide detection band centered at 1300, 1575 and 1700 nm. Fig. SF7(c) directly compares each of the three coefficients separately within the detection bands centered at 1300, 1575 and 1700 nm. Scattering coefficients in the 1575 and 1700 nm bands are consistently lower than those in the 1300 nm band. In contrast, the absorption coefficients in the range centered at 1575 nm are higher than those in the 1300 nm band for a majority of the spectral range detected. Despite this greater absorption, the reduction in scattering results in lower total attenuation across much of the 1575 nm band. Of the three spectral ranges considered, the 1700 nm band has the lowest total attenuation, and the ultradeep OCT approach could yield even greater imaging depths in this wavelength region, assuming a similar detection dynamic range.

In addition to the possibly lower total attenuation coefficient over much of the detection range at 1575 nm compared with 1300 nm, several other mechanisms could be contributing to the increased imaging depth at 1575 nm seen in this study. First, previous studies have suggested that a reduction in the scattering coefficient has a relatively greater impact than increased absorption on OCT imaging depth. Although increased absorption reduces the number of backscattered photons that reach the detector, the fraction of photons that remain ballistic or quasi-ballistic increases due to reduced scattering [Sharma, 2008; Kodach, 2010]. Second, the coherence volume at 1575 nm is larger than at 1300 nm, owing to the lower axial resolution when the detection bandwidths are equal. The larger coherence volume allows signal from a greater tissue volume to contribute to the OCT signal, thus improving the SNR [Carrion, 2007; Harper, 2022]. Third, increased water absorption may preferentially attenuate multiply scattered photons owing to their longer path lengths. This preferential attenuation could reduce the multiple scattering background in coherence-gated imaging and improve image contrast, despite the accompanying reduction in desired signal [Kodach, 2010].

**Figure SF7.**
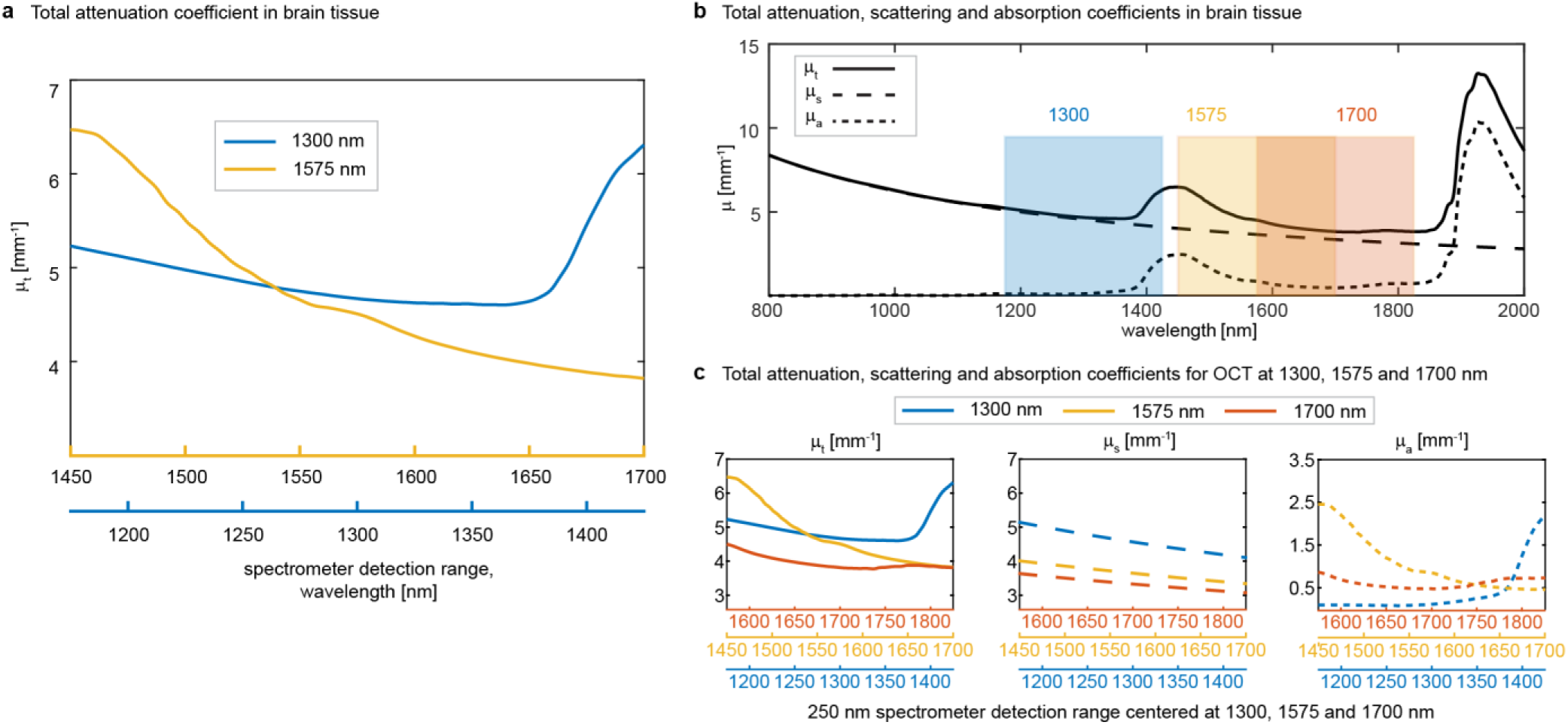
Spectral dependence of calculated attenuation, scattering and absorption in brain tissue. **a,** Total attenuation coefficient, *μ*t, calculated over 250 nm spectrometer detection ranges centered at 1300 and 1575 nm. The two horizontal axes show the corresponding wavelength ranges for the two detection bands. **b,** Total attenuation (*μt*), scattering (*μs*) and absorption (*μa*) coefficients from 800 to 2000 nm. Shaded regions indicate the detection bands centered at 1300, 1575 and 1700 nm. **c,** Direct comparison of *μ*t, *μ*s and *μ*a within the three detection bands. The wavelength axes are offset to align corresponding positions within each band.

We note that there has been one recent study that reported poorer imaging depth at 1550 nm with a custom-built SD-OCT system as compared with a commercial 1300 nm system [Chen, 2025]. However, attribution of this observation to wavelength alone is difficult owing to differences in the system parameters. A similar caveat applies to the present study. Nevertheless, carefully matched experiments have shown increased penetration depth at 1600 nm as compared to 1300 nm [Kodach, 2010] and we present experimental results in biological tissues of interest, along with multiple possible explanations for why greater imaging depths are possible at 1575 nm.

The imaging depths reported in Park et al. [Park, 2018] are most likely optical depths. Their stated values were converted to physical depths, assuming a tissue RI of 1.35. Thus, the reported maximum optical imaging depth of 2.6 mm was interpreted as a physical depth of 1.9 mm. The physical imaging depth of 2.1 mm for Chong et al. [Chong, 2015] was inferred from Figure 3(A) based on the scale bar shown. The structural imaging depth of 2.3 mm, with a 1300 nm swept-source OCT system, reported by Choi et al. [Choi, 2015] was based on intensity profiles and no en-face image at this depth was shown.

### S6: Computational refocusing with bandwidth control

Additional details of the computational refocusing and bandwidth-control procedure are shown in Fig. SF8. The representative B-scan in Fig. SF8(a) was obtained from the same *ex vivo* mouse brain dataset used in Fig. 6. The B-scan is shown after focal-plane flattening, which was performed by reintroducing the estimated coverslip tilt. The focal-plane position was determined using the RMSR method, as described in S2. The focal plane was identified as the position of the minimum in the RMSR profile shown in Fig. SF8(b). The resulting focal-plane estimate is indicated by the pink dashed lines in Fig. SF8(a) and SF8(b). A representative XY plane below the tissue surface and the magnitude of the corresponding transverse spatial-frequency spectrum are shown in Fig. SF8(c). The spatial-frequency spectrum is seen to be displaced from zero spatial frequency [Mulligan, 2019]. The amplitude mask used for bandwidth control had a user-specified normalized radius and a Tukey-tapered edge. Fig. SF8(d) shows the amplitude mask for a normalized radius of 0.9. The mask as well as the refocusing kernel were shifted to align with the center of the measured transverse spatial-frequency spectrum.

Comparison of the coherently averaged images with the results obtained using bandwidth control alone, computational refocusing alone, and computational refocusing with bandwidth control is shown in Fig. SF8(e). At depths of 0 and 0.85 mm, computational refocusing improved the transverse resolution and visibility of selected tissue features, whereas bandwidth control suppressed noise. Combining the two processing steps therefore improved overall feature visibility. At greater depths of 1.7 and 2.55 mm, computational refocusing produced a more limited improvement, while bandwidth control continued to suppress noise and enhance the visibility of weakly backscattering structures. The pink and blue arrows in the first column of Fig. SF8(e) identify representative regions in which feature visibility is improved by computational refocusing and bandwidth control, respectively.

The bright horizontal streaks in the *en face* images at a depth of 0 mm in Figs. 6(a), 8, and SF8(e), which are more prominent before refocusing with bandwidth control, are caused by reflections from the bottom surface of the coverslip. The dark horizo ntal lines in the deepest *en face* images in Figs. 6, 8, and SF8(e) are caused by variations in the AC subtraction and coherent averaging results at the corresponding Y positions.

**Figure SF8.**
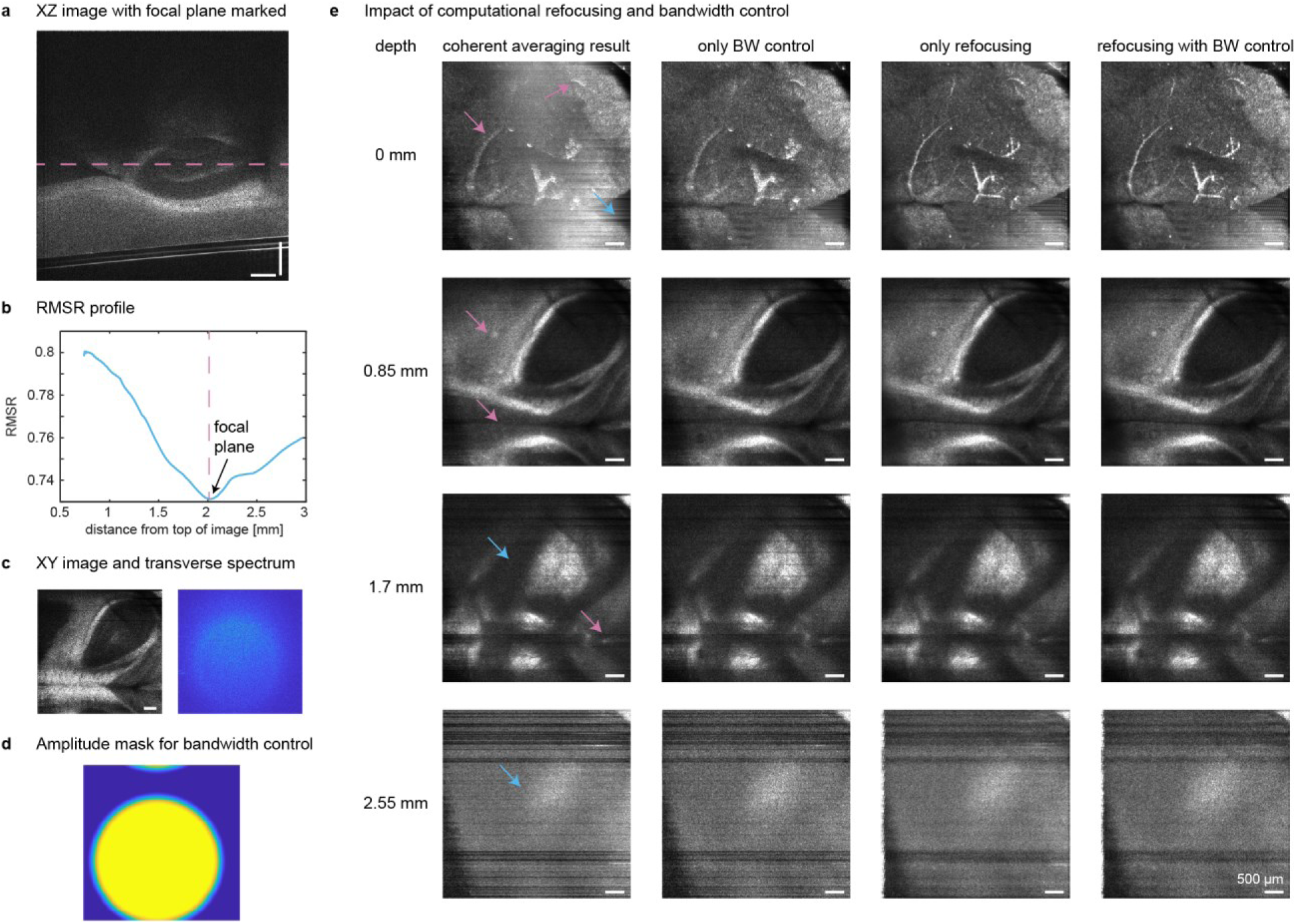
Computational refocusing with bandwidth control. **a,** Representative XZ B-scan of an *ex vivo* mouse brain after focal-plane flattening. The dashed pink line indicates the focal-plane position estimated using the RMSR method. **b,** RMSR as a function of distance from the top of the image. The minimum of the RMSR profile provides the estimated focal-plane position. **c,** Representative XY plane and the magnitude of the corresponding transverse spatial-frequency spectrum. **d,** Amplitude mask used for bandwidth control, shifted to align with the center of the transverse spatial-frequency spectrum. **e,** Effects of computational refocusing and bandwidth control on XY planes located 0, 0.85, 1.7, and 2.55 mm below the tissue surface. From left to right, the columns show the coherently averaged images, images processed using bandwidth control alone, images processed using computational refocusing alone, and images processed using both computational refocusing and bandwidth control. Pink arrows indicate representative regions in which computational refocusing improves feature visibility through increased transverse resolution. Blue arrows indicate representative regions in which bandwidth control suppresses noise and improves the SBR. Scale bars, 500 µm.

### S7: Hippocampal segmentation and volume reconstruction

The features observed in the ultradeep OCT scans were consistent with the expected anatomy of the mouse brain (Fig. SF9). Regional OCT intensity variation attributable to myelin (Fig. SF9(b), SF9(c)), together with the expected shapes and locations of anatomical structures based on the reference atlas in Fig. SF9(d), was used to annotate the enlarged XZ OCT image in Fig. SF9(a). The cerebral cortex exhibited moderate scattering, whereas myelin-rich structures, such as the CC, appeared relatively bright. In contrast, densely packed neuronal layers, such as the pyramidal cell layer of CA1, appeared relatively dark. The dorsal hippocampus was delineated by the surrounding myelin-rich structures and by internal variations in OCT intensity. The stratum lacunosum-moleculare (SLM) of the cornu ammonis (CA) appeared relatively bright and helped distinguish the CA region from the dentate gyrus (DG). There was insufficient contrast to individually delineate CA1, CA2 and CA3 and these outlines in Fig. SF9(a) were drawn by eye. The CA3-hilus boundary was placed based on the expected geometry. Within the DG, the hilus was also distinguishable based on its relatively high OCT intensity. Thin, low-intensity bands were also visible at the expected locations of the pyramidal cell layer of the CA and the granule cell layer of the DG (Fig. SF9(a)). However, these layers did not have sufficient contrast to enable segmentation as independent layers.

The histology section in Fig. SF9(b) is an enlarged version of the SMI-99 immunohistochemistry slice shown in Fig. SF9(c). Fig. SF9(d) is the composite Nissl and annotated atlas section. Histology and atlas images in Fig. SF9(b), SF9(c), SF9(d) are from the Allen Institute mouse brain atlas. Fig. SF9(b) and SF9(c) were derived from the SMI-99 channel data from dataset 100142355, section 128 (image 102167355):https://connectivity.brain-map.org/static/referencedata/experiment/siv/100142355?imageId=102167355&imageType=CALB1&initImage=CALB1. The image in Fig. SF9(d) is atlas image 100960224 [plate 75 of 132]: https://atlas.brain-map.org/atlas?atlas=1#atlas=1&plate=100960224. Histological slices were chosen as representative reference sections and correspond only approximately to the OCT XZ image displayed. The age and sex of the Allen histological sections and reference atlas image (8-week-old, male) are not the same as those of the brains analyzed in this study, which were 32 to 44 weeks old and mostly female.

**Figure SF9.**
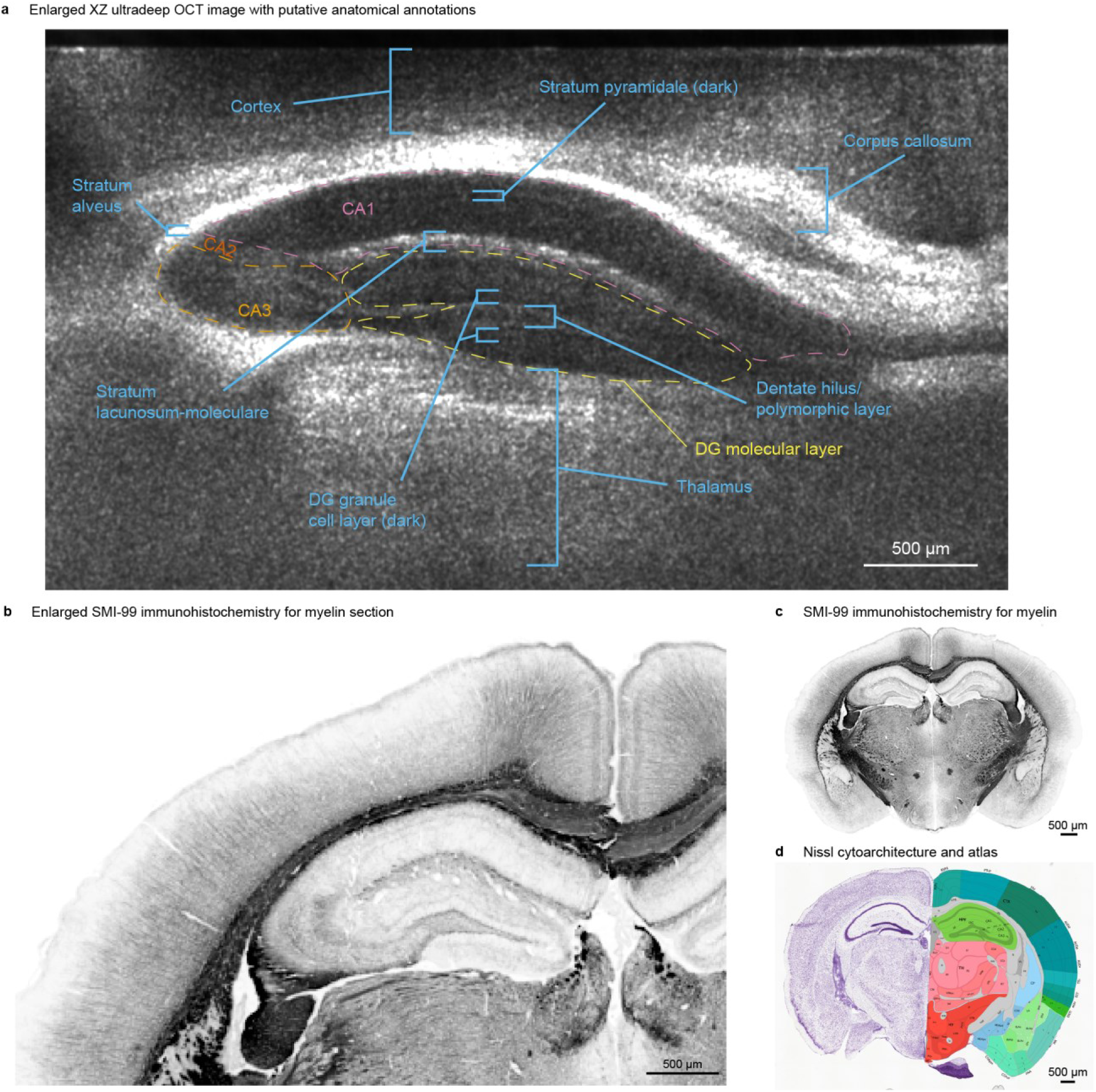
Anatomical interpretation of ultradeep OCT XZ image of the mouse hippocampal region. **a,** Enlarged XZ OCT image with anatomical annotations based on regional geometry and relative OCT intensity. Myelin-rich structures, including the corpus callosum and stratum alveus, appear bright, whereas cell-dense hippocampal layers, including the CA1 pyramidal cell layer and dentate gyrus granule cell layer, appear darker than adjacent tissue. Dashed lines mark hippocampal layers and subfields drawn by eye from atlas geometry and OCT intensity. Boundaries are approximate and not histologically validated. **b,c,** Coronal reference histology section from a male, 56-day-old C57BL/6J mouse, labeled for calbindin (Calb1) and myelin basic protein (SMI-99 antibody). Only the SMI-99 channel is shown, converted to grayscale and inverted; **b** is an enlarged portion of **c**. Adapted from the Allen Mouse Brain Connectivity Atlas reference data [Oh, 2014; Allen Institute, 2011]. Image credit: Allen Institute. **d,** Coronal plate from the Allen Reference Atlas (P56 coronal atlas, male, C57BL/6J reference specimen) showing Nissl-stained tissue in the left hemisphere and the color-coded anatomical annotation in the right hemisphere [Allen Institute, 2011; Dong, 2008]. Image credit: Allen Institute. Histological panels are reference sections and are not registered to the OCT volume. Scale bars, 500 µm.

The segmentation and three-dimensional reconstruction of the hippocampus and its two principal subregions of CA and DG were shown in Fig. 7. Here, we present exploratory segmentations of the same volumes, into four compartments. The dorsal edge of the SLM within the CA was used to split this subregion into CA’ and SLM. The CA’ volume in this case probably included the stratum oriens, pyramidal cell layer and stratum radiatum of the CA1, CA2 and CA3. The SLM compartment likely included the stratum lacunosum-moleculare associated with the CA regions. The DG was split into DG’ and hilus based on the relatively bright appearance of the hilus within the dentate gyrus. The DG’ compartment likely contained the molecular layer and the granule cell layer of the dentate gyrus. The relatively bright appearance of the SLM and the dentate hilus on the OCT images was consistent with the SMI-99 immunohistochemistry shown in Fig. SF9(b), which indicated greater myelin-associated staining in these regions relative to the adjacent hippocampal layers.

The volumetric segmentations of the dorsal hippocampus divided into four compartments: CA’, SLM, DG’, and hilus, are shown in Fig. SF10. Example 1 is the OCT volume from the Alzheimer’s disease model mouse brain (AD) and Example 2 is from the wild-type mouse brain (WT1). The CA volume in Fig. 7 is the union of the CA’ and SLM volumes shown here. Similarly, DG in the main text is a union of DG’ and hilus shown in Fig. SF10. The spatial organization of the four segmented compartments is shown in different views and orientations in Fig. SF10(a). The segmentation contours overlaid on orthogonal XY, YZ, and XZ OCT views show how the compartment boundaries correspond to tissue features across the three planes. In particular, the relatively bright SLM separates the cornu ammonis compartment from the dentate gyrus, while differences in OCT intensity within the dentate gyrus distinguish the hilus from the surrounding molecular and granule cell layers. These orthogonal overlays were used to assess continuity of the anatomical structures on different planes of the OCT volume.

**Figure SF10.**
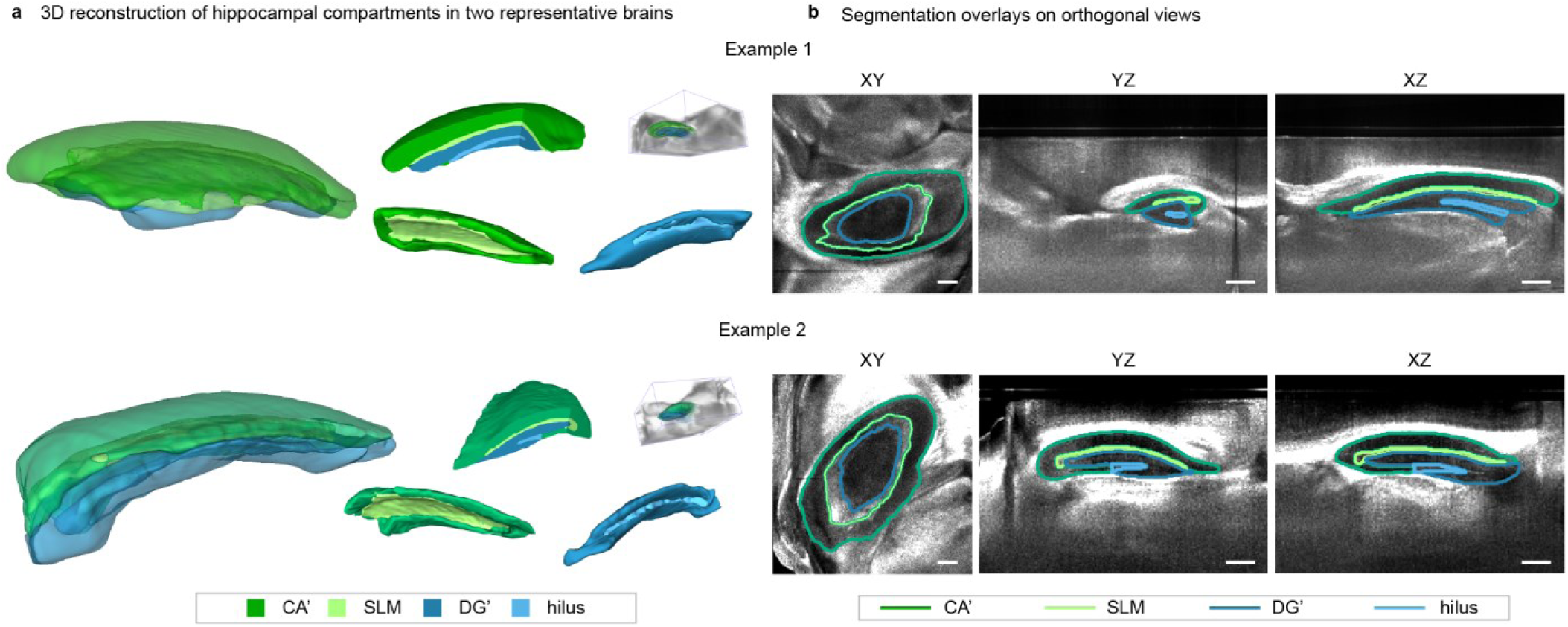
Four compartment reconstructions of the hippocampal volumes shown in Fig. 7. **a,** Reconstructions from Example 1 and Example 2 datasets after segmenting the dorsal hippocampus into four compartments. The views show the entire segmented hippocampus with translucent compartments (left) and cutaway views (top row, middle). Segmented hippocampal volumes within the OCT volumes of dimensions 5 × 5 × 3.8 mm, X × Y × Z are shown on the right. The bottom row displays reconstructions with only the CA’ and SLM compartments in the middle and the DG’ and hilus on the right. CA’, green; SLM, light green; DG’, dark blue; and hilus, light blue. See text for exact compartment descriptions. The 3D renderings are shown in perspective and carry no scale bar; see the orthogonal views in **b** for scale. **b,** Segmentation contours for the CA’, SLM, DG’, and hilus compartments overlaid on representative orthogonal XY, YZ, and XZ OCT views. Scale bars, 500 µm.

